# Human liver organoids; a patient-derived primary model for HBV Infection and Related Hepatocellular Carcinoma

**DOI:** 10.1101/568147

**Authors:** Elisa De Crignis, Shahla Romal, Fabrizia Carofiglio, Panagiotis Moulos, Monique M.A. Verstegen, Mir Mubashir Khalid, Farzin Pourfarzad, Shringar Rao, Ameneh Bazrafshan, Christina Koutsothanassis, Helmuth Gehart, Tsung Wai Kan, Robert-Jan Palstra, Charles Boucher, Jan M.N. IJzermans, Meritxell Huch, Sylvia F. Boj, Robert Vries, Hans Clevers, Luc van der Laan, Pantelis Hatzis, Tokameh Mahmoudi

## Abstract

The molecular events that drive Hepatitis B virus (HBV)-mediated transformation and tumorigenesis have remained largely unclear, due to the absence of a relevant primary model system. Here we propose the use of human liver organoids as a platform for modeling HBV infection and related tumorigenesis. We first describe a primary ex vivo HBV-infection model derived from healthy donor liver organoids after challenge with recombinant virus or HBV-infected patient serum. HBV infected organoids produced cccDNA, expressed intracellular HBV RNA and proteins, and produced infectious HBV. This ex vivo HBV infected primary differentiated hepatocyte organoid platform was amenable to drug screening for both anti-HBV activity as well as for drug-induced toxicity. We also studied HBV replication in transgenically modified organoids; liver organoids exogenously overexpressing the HBV receptor NTCP by lentiviral transduction were not more susceptible to HBV, suggesting the necessity for additional host factors for efficient infection. We also generated transgenic organoids harboring integrated HBV, representing a long-term culture system also suitable for viral production and the study of HBV transcription. Finally, we generated HBV-infected patient-derived liver organoids from non-tumor cirrhotic tissue of explants from liver transplant patients. Interestingly, transcriptomic analysis of patient-derived liver organoids indicated the presence of an aberrant early cancer gene signature, which clustered with the HCC cohort on the TCGA LIHC dataset and away from healthy liver tissue, and may provide invaluable novel biomarkers for disease surveillance and development of HCC in HBV infected patients.

## BACKGROUND

Persistent HBV infection is the leading cause of chronic liver cirrhosis and hepatocellular carcinoma (HCC) world-wide (MacLachlan, 2015; Di Bisceglie, 2009; An, 2018). A combination of viral and host factors determines whether an individual infected with HBV will be able to clear the infection or will become a chronic carrier. Characterized by its high host-species and organ-specificity, HBV infection and replication is thought to orchestrate an interplay between the immune system and viral-specific factors that eventually lead to the onset of HCC. Insights into the molecular mechanisms underlying HBV-induced HCC have largely been provided by epidemiological studies (El-Serag, 2012; Fattovich, Bortolotti, & Donato, 2008; Jiang et al., 2012; Sagnelli, Macera, Russo, Coppola, & Sagnelli, 2019), genome wide analysis of viral and host characteristics (Cancer Genome Atlas Research Network. Electronic address & Cancer Genome Atlas Research, 2017; Fujimoto et al., 2012; Huang et al., 2012; Ji et al., 2014; Sartorius et al., 2019; Shibata & Aburatani, 2014; Sung et al., 2012)), as well as by studies performed in in vitro settings using hepatoma cell lines (Thomas & Liang, 2016; Zhang, Wang, & Ye, 2014).

However, a major deficiency attributed to the strict viral host and cell type tropism is the limited availability of relevant animal or in vitro model systems to study HBV infection. Chimpanzees remain the only animal model that supports the full HBV replication cycle, while available hepatoma cell line models are unsuitable for delineating the molecular steps towards tumorigenesis as they differ substantially from primary cells in their already tumor-derived gene expression profiles (Protzer, 2017). These systems have inherent limitations resulting in poor predictive value for clinical outcome (Allweiss & Dandri, 2016). Primary hepatocytes present the gold standard model system for HBV research in vitro, especially with recent studies that have demonstrated a significant increase in the half-life of these cultures (Xiang et al., 2019). However they are difficult to obtain and, since they cannot be expanded, are not usually available in quantities sufficient to perform large-scale analyses (Hu, 2019). Induced pluripotent stem cell (iPSC)-derived hepatocytes which are susceptible to HBV infection and support replication are also a useful model for studying host-virus determinants of replication (Kaneko et al., 2016; Nie et al., 2018; Sakurai et al., 2017; Xia et al., 2017). However, iPSC-derived hepatocyte models cannot be patient derived, limiting studies to only ex-vivo infection systems and limiting the possibility of patient-specific personalized treatment approaches (Torresi, 2019; Nantasanti, 2016). As a consequence of deficiencies in available model systems, despite its fascinating biology, many questions regarding the life cycle of HBV and its mechanisms of persistence, including HBV-induced molecular events underlying tumorigenesis, remain largely unexplored in primary settings, and key viral and host players involved remain unknown.

The dependence of hepatocytes on spacial and matrix-derived signals had until recently prevented their long-term in vitro culturing. The organoid culture technology involves the generation of cell-derived genetically-stable in vitro 3D-organ models of human origin. We have previously established a primary liver culture system based on isolation and expansion of primary cells that allows for the long-term expansion of liver cells as organoids (Huch et al., 2013; Huch et al., 2015). In this culture system, isolated adult hepatic cells are expanded through multiple passages in an optimized medium (EM) without induction of genomic alterations (Huch et al., 2015). When switched to differentiation medium (DM) where proliferation signals are removed and ductal (progenitor) fate is inhibited, liver organoid cultures differentiate into functional hepatocytes in vitro as exemplified by their polygonal cell shape (Figure 1a), and hepatocyte functions including Albumin production and Cytochrome C3A4 expression and activity (Huch et al., 2015). Here we use the human liver organoid platform to model and study HBV infection and replication, as well as related tumorigenesis in patient-derived organoids generated from HBV-infected donors. This expandable model yields patient-specific organoids in quantities amenable to molecular and functional characterization, and allowed us to generate a living biobank of HBV infected patient-derived cells amenable to downstream genomic, transcriptomic and proteomic analysis as well as screening for HBV-directed therapeutics. We first describe the ex-vivo HBV-infection of healthy donor (hD)-derived liver organoids, as a model to investigate the viral infection and replication in hepatocytes. We use the HBV infected organoid model as a platform for drug screening that can measure both drug-induced anti-HBV transcription and replication activity as well as drug-induced toxicity. We also demonstrate that transgenic modification of liver organoids provides an in vitro mechanistic platform to study the molecular determinants of HBV infection and replication. Finally, we conduct transcriptomic analysis of HBV-infected patient-derived organoids and describe the discovery of an early cancer gene signature, a potentially invaluable prognostic biomarker for HCC.

**Fig. 1:**
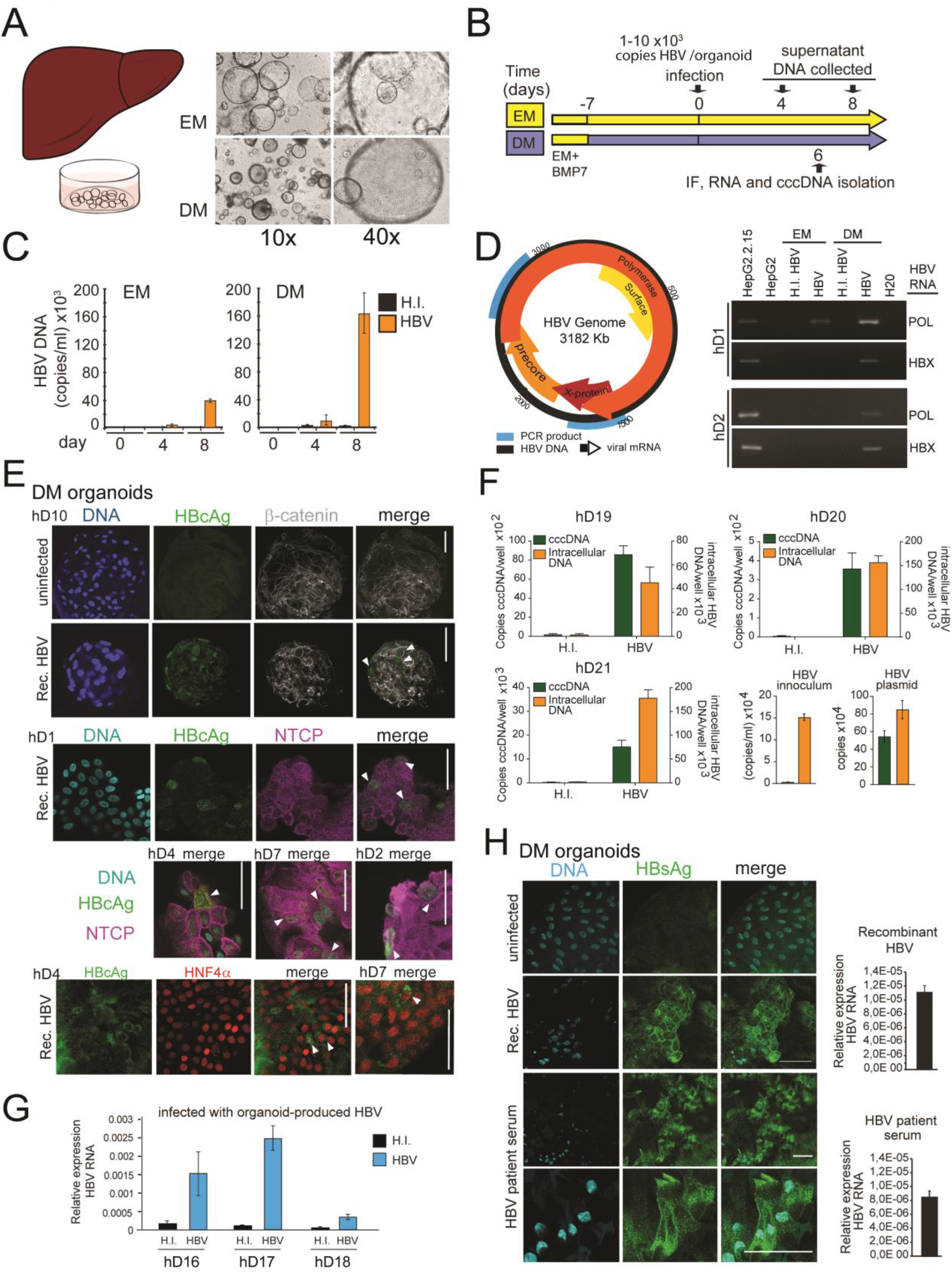
Modeling HBV infection in vitro using human liver organoids. (A) Representative images of liver organoids in EM and DM (B) Experimental design of infection experiments. Arrows indicate the time points for HBV detection. (C) Levels of HBV DNA in supernatant of infected organoid cultures were quantified at indicated times by real time PCR and compared to the cultures challenged with HI virus. (D) Schematic of the HBV genome showing ORFs (arrows) and the localization of PCR products (blue boxes). The agarose gel demonstrates expression of HBX and POL genes by nested PCR performed on cDNA obtained from two in vitro infected healthy donor (hD) organoid lines. (E) Immunofluorescent staining showing the expression of HBcAg (green) together with NTCP (magenta), β-catenin (grey) or HNF4α (red) performed in different hD organoids 6 days after HBV infection in differentiation medium. (F) Quantification of total HBV DNA (orange) and cccDNA (green) from intracellular DNA purified from three healthy donors (hD) organoid lines 6 days post infection with HBV under DM condition. Quantification of total HBV DNA and cccDNA is also shown from supernatant of infected organoids (inoculum) and from double stranded HBV plasmid (as positive control for cccDNA) (G) Expression of intracellular HBV RNA relative to beta 2 microglobulin in three hD organoid lines infected with organoid-produced HBV (concentrated from pooled supernatants of organoid cultures infected with HepG2.2.15-produced HBV). (H) Immunofluorescent staining showing the expression of HBsAg (green) in DM organoids infected with recombinant HBV and patient serum. Scale bars represent 50 μm. Bar graphs show total HBV RNA levels in the culture at the time of staining.

## Results

### Human liver organoids allow modelling of HBV infection in vitro

We first used the previously characterized liver organoid platform (Huch et al., 2015) generated from healthy donors to set up a novel ex-vivo HBV-infection system to study HBV replication. Liver organoids from healthy donors were grown in either expansion media (EM) or differentiation media (DM) (Figure 1a) for 7 days prior to infection with recombinant HBV generated from HepG2.2.15, a HepG2 cell line subclone stably expressing HBV (Figure 1b-c). As control for the inoculum, we also infected organoids with heat inactivated HBV (HI). HBV infection and replication were validated by quantifying the levels of HBV DNA in the supernatant (Figure 1c), intracellular HBV RNA (Figure 1d), HBV-specific proteins by immunofluorescence microscopy (Figure 1e) and intracellular covalently closed circular DNA (cccDNA) from infected organoids (Figure 1f). HBV DNA was detected in organoid culture supernatants from four days post infection, but not from the heat-inactivated virus infected cells, indicating successful HBV replication (Figure 1c). Differentiated organoids maintained in DM were more efficiently infected and produced higher viral titers than organoids maintained in EM (Figure 1c-d). The RNA intermediates necessary for protein production and viral replication (HBV pol and Hbx) were present in infected DM organoids and detected by RT-PCR analysis, but not in the heat-inactivated virus infected cells (Figure 1d). Immunostaining, using antibodies recognizing HBV core (HBcAg) showed specific nuclear and cytoplasmic staining in multiple infected healthy donor liver organoid lines, confirming the presence of foci of HBV replication in HBV-infected cells predominantly in infected DM organoids (Figure 1e and Figure 1-Figure Supplement 1). Furthermore, infection of DM organoids resulted in the production of cccDNA, a definitive marker of HBV replication, as detected by a qPCR-based cccDNA detection method of intracellular HBV DNA after digestion with a nuclease to specifically remove non-cccDNA (Figure 1f). Inoculum that lacks cccDNA was used as a negative control for the cccDNA-specific qPCR and HBV plasmid DNA was used as a positive control (Figure 1f). HBV replication, infection and spread appeared to be persistent until 8 days after infection when viral production dropped significantly, likely because of the limited half-life of differentiated organoids in culture (Figure 1-Figure Supplement 2). Periodic culturing of the organoids in EM in order to stimulate the recovery and proliferation of the organoids modestly extended the half-life of the infected cultures, where viral production was maintained for approximately one month post infection (Figure 1-figure supplement 2). To determine whether the organoids are capable of producing infectious HBV, supernatants containing virus produced by organoids, were collected, concentrated and used for subsequent spinoculation of healthy donor (hD) organoids. As shown in Figure 1g, infection of hD organoids with organoid-produced HBV resulted in expression of intracellular HBV RNA, indicating that organoids produce infectious viral particles. Growth of viral isolates from patient material has been limited by the lack of an adequate primary model system. However, differentiated organoids were able to support infection and replication when challenged with HBV infected patient sera, as shown by production of viral DNA, expression of viral transcripts and positive immunostaining for HBsAg (Figure 1h and (Figure 1-figure supplement 3). Differentiated liver organoids therefore provide a useful ex vivo HBV infection platform in which the role of specific host and viral factors can be investigate.

### Ex vivo HBV-infected liver organoids are a viable platform for anti-viral drug screening and drug induced toxicity

We next examined whether the ex vivo infected liver organoid platform would be amenable to anti-HBV drug screening to monitor antiviral activity and drug-induced toxicity of two different drugs, Tenofovir and Fialuridine, according to the schematic in Figure 2a. Tenofovir is a nucleoside reverse transcriptase inhibitor which inhibits the reverse transcription of HBV pre-genomic RNA to DNA. Fialuridine, also a nucleoside analogue that inhibits reverse transcription, was shown to cause severe hepatotoxicity in patients (McKenzie et al., 1995). In the organoids, HBV viral DNA production in the culture supernatant was inhibited by both Tenofovir and Fialuridine in three independent healthy donor (hD) derived organoids; whereas, as expected, RNA levels remained the same (Figure 2b). Therefore, the organoid ex vivo-infection platform not only allows measurement of drug-induced antiviral activity, but also offers insight into the mechanism of drug action by allowing delineation of which step of the HBV life cycle is targeted and inhibited. As expected, treatment of HepG2.2.15 cells with Tenofovir and Fialuridine resulted in similar decreases in released HBV DNA, but no change in intracellular HBV RNA levels (Figure 2c), reaffirming the mechanism of action of these drugs in a cell-line model of HBV replication. Due to the well-established detrimental effects of Fialuridine on viability of primary human hepatocytes, we sought to evaluate Fialuridine-induced toxicity on primary human liver organoids as well as in HepG2 cells. We measured the viability of organoids and HepG2 cells using the alamarBlue viability assay and by monitoring their phenotype upon Fialuridine and Tenofovir treatment using microscopy. HepG2 cells demonstrated no change in cell viability upon treatment with Tenofovir and increasing concentrations of Fialuridine as compared to the mock treated cells (Figure 2d). The phenotype of HepG2 cells was also comparable across all treatments as observed by microscopy (Figure 2e). Strikingly, the liver organoids treated with Fialuridine at as low a concentration of 1μM demonstrated a significant reduction in viability as measured by AlamarBlue assay when compared to mock treated cells (Figure 2f and Figure 2-figure supplement 1a). The organoids treated with higher Fialuridine concentrations (5-20μM) as well as with 20μM Tenofovir also demonstrated impaired viability (Figure 2f and Figure 2-figure supplement 1a). The decreased cell viability was also apparent in the phenotype of the 1-20μM Fialuridine-treated and 20μM Tenofovir-treated organoids compared to vehicle control as observed by microscopy (Figure 2g and Figure 2-figure supplement 1b). This highlights that Fialuridine-induced toxicity is evident and quantifiable in the primary human liver organoid model but not in the HepG2.2.15 model of HBV replication. Thus, we demonstrate that ex vivo infected differentiated liver organoids support the full replication cycle of HBV and serve as an ideal novel primary platform for drug screening and to elucidate the molecular events underlying HBV replication. Moreover, human liver organoids serve as an ideal platform for monitoring drug induced toxicity in pre-clinical studies.

**Fig. 2:**
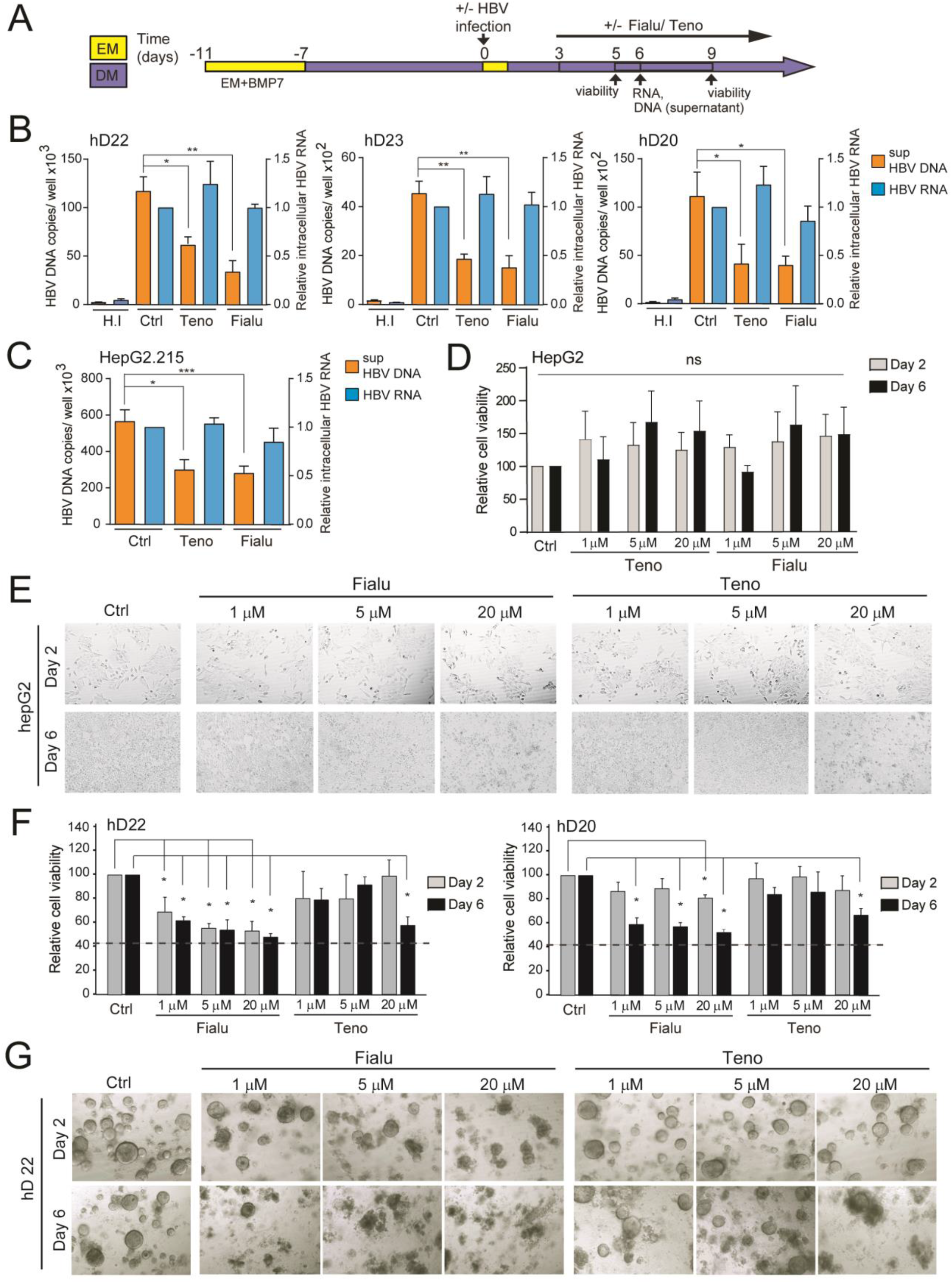
HBV infected liver organoids as a model for HBV antiviral drug screening and toxicity. (A) Experimental design of drug treatment of HBV infected liver organoids followed by assessment of antiviral activity and toxicity. Arrows indicate time points for HBV detection or assessment of viability. Levels of HBV DNA in the supernatant and intracellular HBV RNA were quantified by real time PCR and RT-PCR respectively for three independent healthy donors (B) and HepG2.2.15 cells (C) upon treatment with control vehicle, Fialuridine (10μM) or Tenofovir (10μM) as indicated. Data are shown as mean ± SD of at least 3 replicate treatments \**P* < 0.05; \*\**P* < 0.01. (D) Relative viability of HepG2 cells was measured using the AlamarBlue cell viability assay after treatment with vehicle control, Fialuridine or Tenofovir for 2 or 6 days as indicated, normalized to vehicle control and plotted as the average of percent viability ± SD (n=3) (n.s = not significant) (E) Representative bright field images taken of HepG2 cells treated with antiviral drugs for 2 or 6 days as indicated. (F) Bar diagrams representing relative cellular viability of hD liver organoids after 2 or 6 days of treatment with Fialuridine or Tenofovir at the different concentrations indicated using the AlamarBlue cell viability assay. All values are normalized to the vehicle treated control and plotted as the average of percent viability ± SD (n=3). The dotted line represents the lower limit of quantification based on values obtained from wells free of organoids containing the BME matrix only. (G) Representative bright field images taken of liver organoids treated for 2 or 6 days with the vehicle control or increasing concentrations of the antiviral drugs Tenofovir or Fialuridine as indicated.

### HBV replication can be investigated in transgenically-modified liver organoids

We previously observed higher HBV infection efficiency in DM organoids as compared to EM organoids (Figure 1c). This correlated with higher levels of Sodium taurocholate co-transporting polypeptide (NTCP) expression, a cellular receptor expressed on the surface of hepatocytes implicated in HBV entry (Yan et al., 2012), in differentiated organoids as compared to organoids in EM (Figure 3a-b). Exogenous expression of NTCP in hepatoma cell lines was shown to confer susceptibility to infection (Yan et al., 2012) in line with our observed increased HBV infection in differentiated organoids, likely because of the higher level of NTCP expression. Since differentiated organoid cultures have a limited half-life, we sought to generate transgenically modified healthy donor (hD) organoids exogenously expressing NTCP under expansion conditions (Figure 3c-e and Figure 3-figure supplement 1a-b) in order to improve infection efficiency and facilitate downstream analyses and investigation of the molecular events involved in HBV replication. We used a lentiviral construct harboring the coding sequence of Flag-tagged NTCP ubiquitously expressed under a CMV promoter, followed by a blasticidin selection marker (Figure 3c). Immunofluorescence experiments performed on NTCP-liver organoids in expansion phase confirmed high levels of NTCP protein expression correctly localized to the cellular membrane (Figure 3e and Figure 3-figure supplement 1b). Cholesterol target genes were induced in response to statin treatment in NTCP transgenic organoids, confirming the functionality of exogenously expressed NTCP (Figure 3-figure supplement 1c). We then evaluated viral production following HBV infection in transgenically modified-NTCP organoid lines as compared to parental lines (Figure 3f-g). Interestingly, comparable levels of HBV DNA and HBS antigen levels were observed in the supernatants of both parental and NTCP expressing organoid lines (Figure 3f-g), suggesting that expression of NTCP alone is not sufficient to improve HBV infection rate in liver organoids grown in EM (Figure 3f-g and Figure 3-figure supplement 1d).

**Fig. 3:**
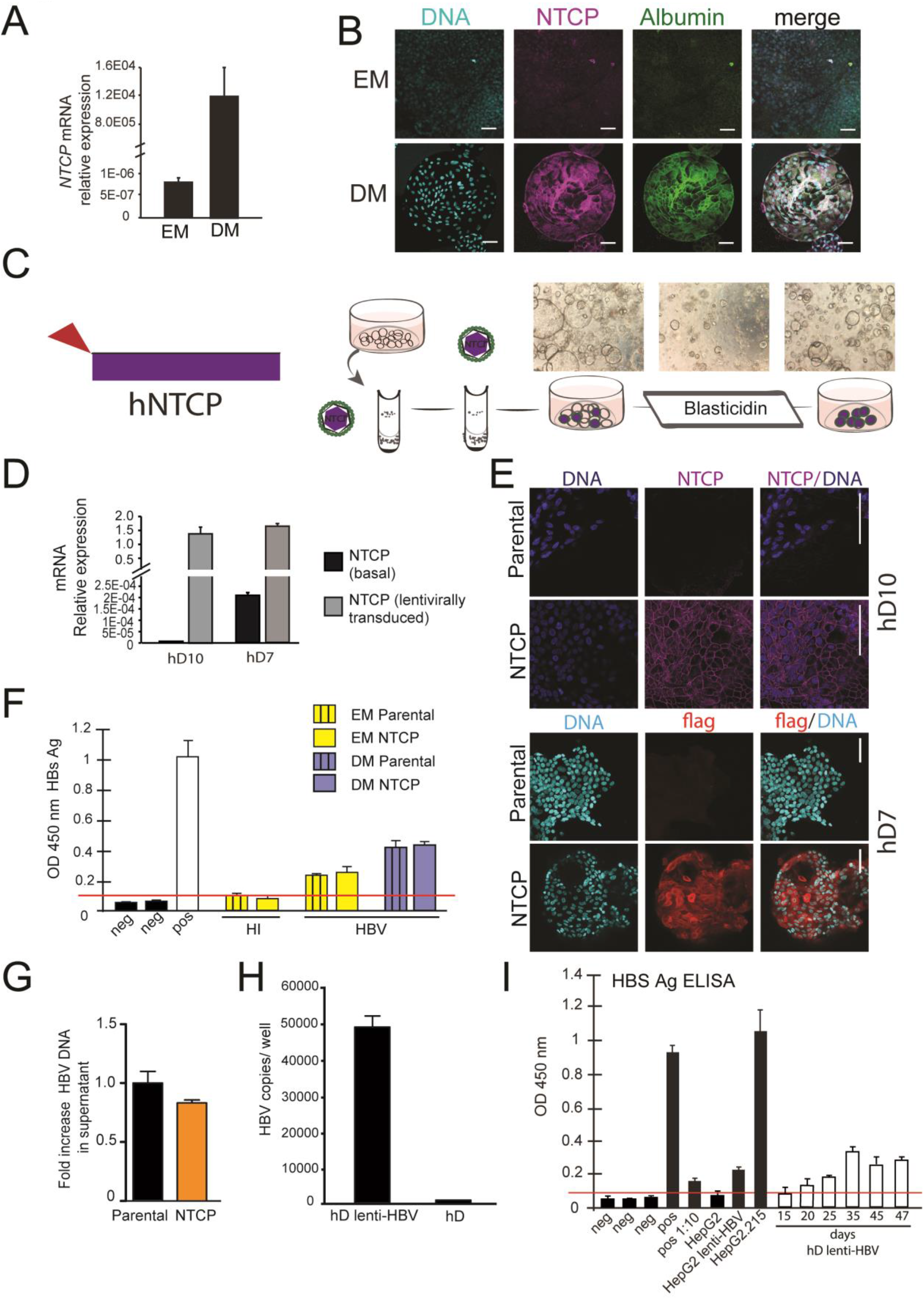
Lentivirally transduced transgenic liver organoids in the study of HBV. (A) mRNA expression levels of the HBV receptor NTCP in undifferentiated (EM) and differentiated (DM) organoids (n=3); mRNA levels were calculated according to the 2ΔCt method using GAPDH as reference gene. (B) Immunofluorescent staining showing the expression of NTCP (magenta) in EM and DM organoids. Nuclei were counterstained with Hoechst 33343 (cyan). Scale bars represent 50 μm. (C) Schematic representation of the experimental procedure for the transduction experiments. Following infection with a lentiviral vector expressing Flag-NTCP, organoids were selected with blasticidin for 5 days in order to obtain lines expressing NTCP in the expansion phase. (D) Levels of expression of NTCP were evaluated by RT-PCR in the untransduced (Parental) and the transduced (NTCP) lines. Expression of NTCP was calculated according to the 2ΔCt method using the housekeeping gene GAPDH as reference gene and confirmed by immunofluorescence staining targeting NTCP (magenta) or Flag (red) (E). (F) HBsAg released in the supernatant of parental and NTCP organoid lines grown in EM or DM 10 days after HBV infection was detected by ELISA. Challenge with heat inactivated virus was used to control for HBsAg present in the inoculum. Pos and neg bars correspond to positive and negative controls provided by the kit manufacturer. Threshold for positivity (red line) was calculated as the average OD + 2SD of negative controls. (G) HBV DNA in the supernatant of NTCP expressing organoid cultures was quantified 5 days after infection and compared to DNA detected in the supernatant of untransduced HBV infected organoids (n=3). Bars represent fold increase in HBV DNA detected in the supernatant, untransduced HBV infected organoids were used as reference. Relative amounts of (H) HBV RNA and (I) HbS antigen produced by hD lenti-HBV organoid lines.

To further highlight the ability of liver organoids to be transgenically modified, we produced a long-term, expandable, primary HBV producing liver organoid model system that can be used to study HBV transcription events. We utilised a lentiviral construct to produce transgenic organoid lines containing an integrated copy of HBV (Figure 3-figure supplement 2). Interestingly, when the HBV genome was placed downstream of the CMV promoter (Figure 3-figure supplement 2c), transgenic organoids, while efficiently expressing HBV RNA, did not support replication, as demonstrated by the low levels of HBV DNA in the culture supernatant (Figure 3-figure supplement 2d), pointing to CMV promoter-induced transcriptional interference with promoter usage required for mRNA transcription of individual HBV genes. Removal of the CMV promoter to generate “lenti-HBV” (Figure 3-figure supplement 2e-f) allowed for the generation of replication-competent, expandable, long term, hD transgenic lenti-HBV organoid lines, in which transcription from the HBV transgene results in protein and viral production (Figure 3h-i, Figure 3-figure supplement 2f). Although lacking cccDNA, this model system provides a primary platform to screen for inhibitors of HBV transcription and is a primary-cell alternative to the HepG2.2.15 cell lines which also do not produce cccDNA, for studies into HBV pathogenesis. Thus, the amenability of human liver organoids to transgenic modification enables investigation of HBV replication and in depth characterization of the molecular events involved.

### HBV-infected patient-derived liver organoids contain functional hepatocytes

The ability to generate a patient-derived primary model is a key advantage of using the liver organoid platform. We applied the previously characterized method to generate liver organoids from healthy donors (Huch et al., 2015) to generate novel patent-derived organoids from HBV-infected individuals undergoing liver transplantation (Figure 4a). The explant used for generating patient-derived organoids was HBV-infected, chronically cirrhotic liver tissue obtained from eleven explanted livers from people infected with HBV (Figure 4a and b, Table 1). We generated and expanded organoid cultures from fresh and frozen explant tissue from all donors with similar efficiency (data not shown).

**Fig. 4:**
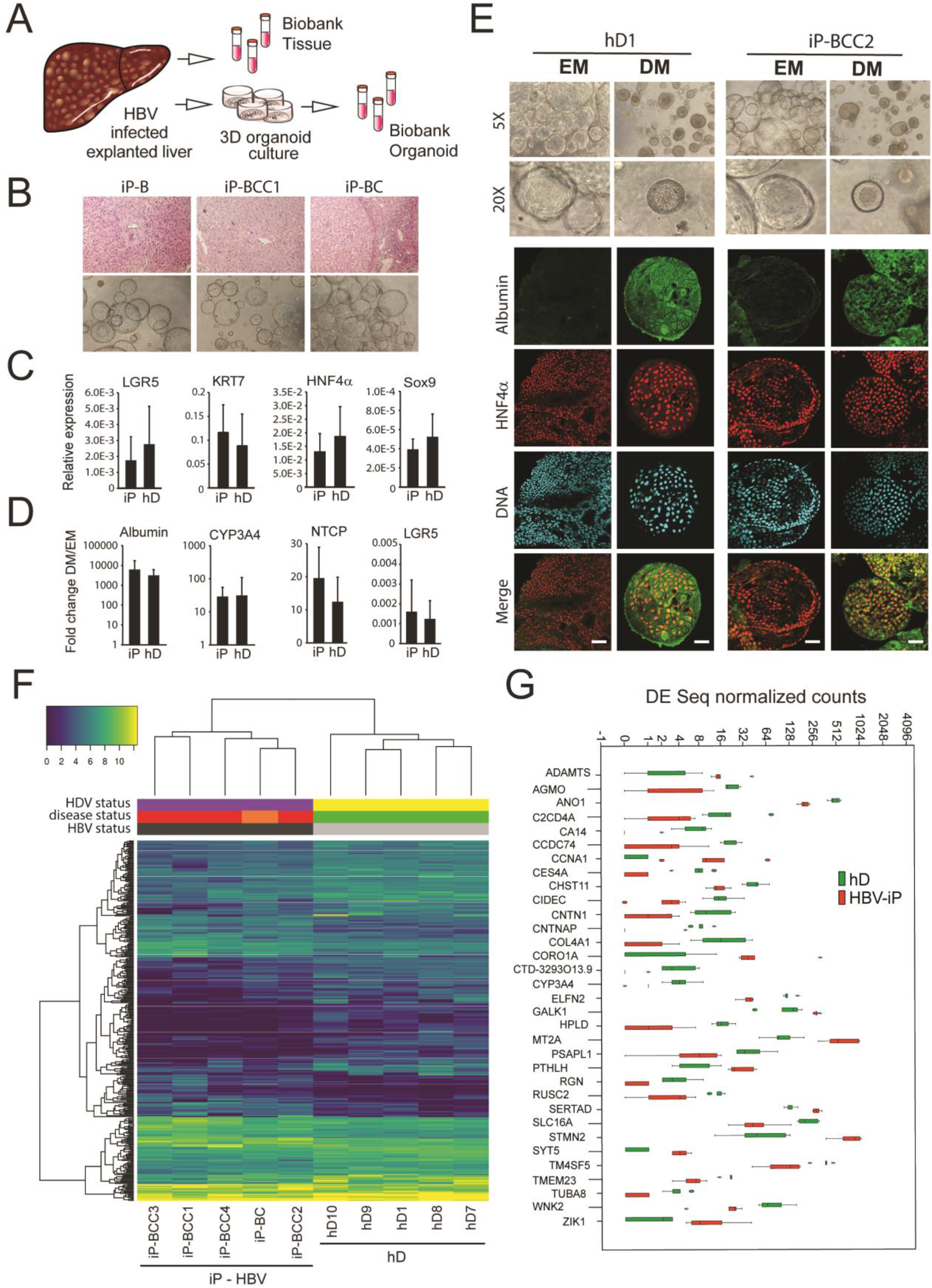
Characterization of organoid cultures from liver explants of HBV-infected patients. (A) Representative panel showing the procedure to generate organoid cultures or biobanks from liver tissue. (B) Hematoxylin-eosin stained sections of explanted liver tissue and phase contrast pictures showing the morphology of liver organoids derived from HBV infected individuals. (C) Expression profile of the progenitor markers LGR5, KRT7, HNF4α, and Sox9 in EM (undifferentiated) organoids derived from liver of healthy donors (hD) (n=4) and HBV infected individuals (iP) (n=5). Levels of expression were calculated according to the 2ΔCT method using GAPDH as reference gene (D) Differentiation capacity of organoid cultures derived from liver of hDs (n=4) and iPs (n=5). Bars represents fold difference in the expression of hepatocyte-specific genes Albumin, Cytochrome CYP3A4, NTCP and of the progenitor-specific gene LGR5 in DM (differentiated) cultures compared to EM organoids using the 2ΔΔCT method. (E). Immunofluorescent staining targeting albumin (green) and HNF4α (red) was performed in EM and DM organoids. Phase contrast images, depicting the morphology of the cells are shown as reference. (F) Hierarchical clustering heatmap of differentially expressed genes derived from the comparison between the group of five hDs and five iPs presenting HBV infection and HCC (four out of five). (G) Box plot of DESeq normalized counts of 33 putative biomarker genes obtained from healthy donor organoids (depicted in green) or obtained from HBV-iP organoids (depicted in red).

Non-tumor, HBV-infected, patient-derived (infected patient (iP)) organoids were expanded in culture (EM) and displayed proliferation rates (Figure 4b and data not shown) and expression of progenitor markers LGR5, KRT7, HNF4α, and Sox9 (Figure 4c) comparable to that of healthy donor (hD) organoids. When grown in differentiation medium (DM), in which proliferation signals are removed and the progenitor fate is inhibited, liver organoid cultures from both healthy and HBV-infected sources acquired hepatocyte fate and differentiated into functional hepatocytes, similar to previously obtained data for healthy liver organoids (Huch et al., 2015). In DM, both hD and iP liver organoid cultures (Huch et al., 2015) showed increased expression of hepatocyte-specific genes Albumin and Cytochrome C3A4, and the cellular receptor implicated in HBV hepatocyte entry NTCP, concomitant with decreased expression of the stem cell-specific gene LGR5 (Figure 4d and Supplementary Figure 6). None of the iP organoids showed signs of HBV production at the RNA, DNA or protein level (data not shown), suggesting that adult stem cells from which the liver organoids are generated are not infected by detectable levels of HBV. At the phenotypic level, while EM organoids grew larger in size and were translucent, differentiated iP organoids showed hepatocyte morphology and a thickening of the outer cell layer, comparable to hD organoids (Figure 4e and Figure 4-figure supplement 1a) and produced comparable levels of albumin as detected by immunofluorescence staining (Figure 4e and Figure 4-figure supplement 1b). Thus, non-tumor HBV iP and hD organoids display comparable phenotypes, retain the capacity for differentiation and are conducive to downstream genomic, transcriptomic and proteomic analysis.

### HBV infected patient-derived liver organoids display a distinct early gene expression signature

The early detection of HBV-related HCC is a challenge that remains critical to direct optimal clinical management of the disease. Despite widely practiced periodic surveillance of patients with cirrhosis, patients with HCC are mostly diagnosed in a late stage. The presence of diagnostic biomarkers for early events in liver cell tumorigenesis would therefore be invaluable to early detection. In order to identify potential early biomarker genes for HBV-induced HCC, we performed mRNA sequencing of the organoid lines derived from HBV-infected patient (iP) and compared their gene expression profile to that of organoids derived from healthy donors (hD). We performed hierarchical clustering of protein-coding differentially expressed genes obtained from the comparison between organoids of five hD organoid lines with five iPs (Figure 4f, Table 1). The iP organoid lines were seeded from five HBV mono-infected patients with cirrhotic liver, four of whom presented with small tumors at the time of explant (Figure 4f, Table 1). Interestingly, although the hD and iP organoids were phenotypically indistinguishable, the iP organoids clustered separately from the healthy donors (Figure 4f). This comparison resulted in identification of an iP-characteristic “gene signature” (Figure 4f and Table 2). We then ranked the differentially expressed genes according to the relative distance of their expression in hD versus iP derived organoids and identified a group of 33 putative early biomarker genes (Figure 4g). GO-term and KEGG-pathway analysis revealed that the “early signature” genes were enriched in metabolic pathway-associated genes (Table 3). Among these, CCNA1, STMN2, which we found were upregulated in the non-tumor patient-derived organoids, were previously identified to be upregulated in HCC (Allain, Angenard, Clement, & Coulouarn, 2016; Chen et al., 2019; Gao et al., 2008; Paradis et al., 2003). Conversely, WNK2, RUSC2, CYP3A4 and RGN, among the significantly downregulated genes in the non-tumor iP organoids have been described as tumor suppressors downregulated in HCC (Allain et al., 2016; Ashida et al., 2017; Tao et al., 2011; Yamaguchi, 2015). Therefore, transcriptomic analyses of healthy vs. patient-derived organoids resulted in the identification of an HBV infection early gene signature and possible biomarkers for HBV infection.

### Transcriptomic analysis of HBV-patient derived organoids result in identification of cancer gene signature

We then applied this “early gene signature” derived from our analysis (Figure 4f) to all our liver organoid samples. Our subsequent analysis included 5 healthy donors (hD1-5), 10 HBV-infected samples (iP) and one donor that had previously been infected with HBV, but had subsequently cleared infection and displayed no aberrant phenotype (infected donor iD). Amongst the iPs, there were samples from patients that had HCC at the time of collection (iP-HBV-HCC); acute liver failure (iP-ALF-HBV); an HBV-infected cirrhotic liver without HCC, no acute liver failure or any coinfections (iP-BC); and HBV-HDV coinfected samples that had either HCC (iP-HCC-HBV+HDV) or acute liver failure (iP-ALF-HBV+HDV). Table 1 describes all patients, groups and clinical characteristics. The early signature when applied to all iP and hD samples, perfectly groups the healthy (grey) against the HBV-infected samples (black) (Figure 5a). HBV-infected iP organoids (purple) and the HBV/HDV co-infected iP organoids closely together (light blue). The signature also separates and groups together the non-infected samples (green), iP organoids derived from patients who had HCC (red) as well as those derived from patients with acute liver failure (dark blue). Interestingly, the organoids generated from the infected donor (iD) who had cleared HBV infection (light orange), as well as the infected patient with cirrhotic HBV-infection (dark orange) clustered together and closely to the larger group of iP organoids derived from patients with HCC (red) (Figure 5a). This observation from iD clustering closely with iP-HBV-HCC indicated the presence of the early HCC-like gene signature in this infected donor liver despite clearance of HBV infection and absence of phenotypic and functional abnormalities at time of donation, an observation that may be important to considered for transplantation purposes and surveillance. In agreement, multidimensional scaling analysis of all iP and hD samples indicated that all hD organoids are clearly separated from the rest and tightly grouped together, while the iP organoids form separate groups corresponding to HDV coinfection, HCC, acute liver failure or previous/cirrhotic HBV infection (Figure 5b). We have therefore identified a specific gene signature/group which discriminates the hD and iP organoids among more complex classifications (Figure 5a-b). We applied this gene signature to all samples stored in The Cancer Genome Atlas Liver Hepatocellular Carcinoma (TCGA-LIHC) database, a depository of sequences from HCC patients. The application of this gene signature when mapped onto the relevant TCGA gene expression data distinctly separated the HCC samples from non-HCC tissue (Figure 5c). This important observation indicates that transcriptomic analysis of our patient-derived liver organoid model identified a novel early liver cancer gene signature in non-tumor HBV-infected patient-derived liver organoids, despite the absence of phenotypic signs of aberrant growth. Thus, HBV-infected patient-derived liver organoids are a novel primary 3D cell culture model that resemble the diseased tissue of origin and can be used from genomic, transcriptomic, proteomic and clinical applications to identify biomarkers for disease states during HBV infection.

**Fig. 5:**
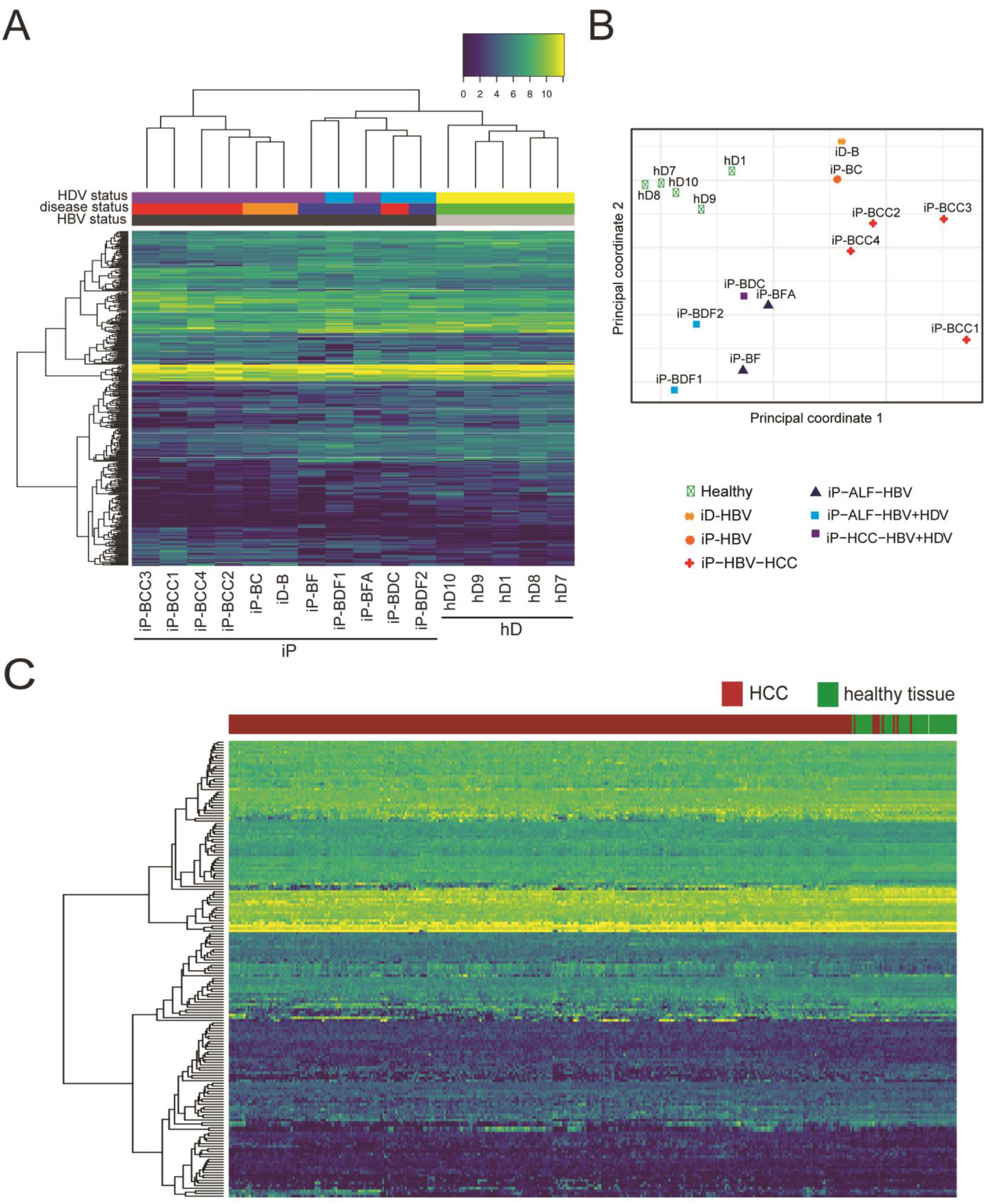
Identification of early HCC gene signature in non-tumor iP-derived organoids. (A) Hierarchical clustering heatmap of all iP and hD samples depicting the grouping and expression levels of protein coding differentially expressed genes derived from the comparison between the group of five hDs and five iPs presenting HBV infection and HCC (four out of five). Coloured bars on top of the heatmap indicate HBV status (black for HBV positive patients, gray for healthy donors), disease status (red for HBV positive HCC, orange for HBV positive without HCC, blue for HBV positive acute liver failure, green for healthy donors), and HDV coinfection status (purple HBV positive HDV negative, light blue for HBV positive HDV positive, yellow for healthy donors). (B) Multidimensional scaling plot of all iP and hD samples. (C) Hierarchical clustering heatmap of liver HCC gene expression data from 342 HCC tissue samples and 47 samples from matched nearby tissue from TCGA using the HBV iP versus hD gene signature.

**Fig. 6:**
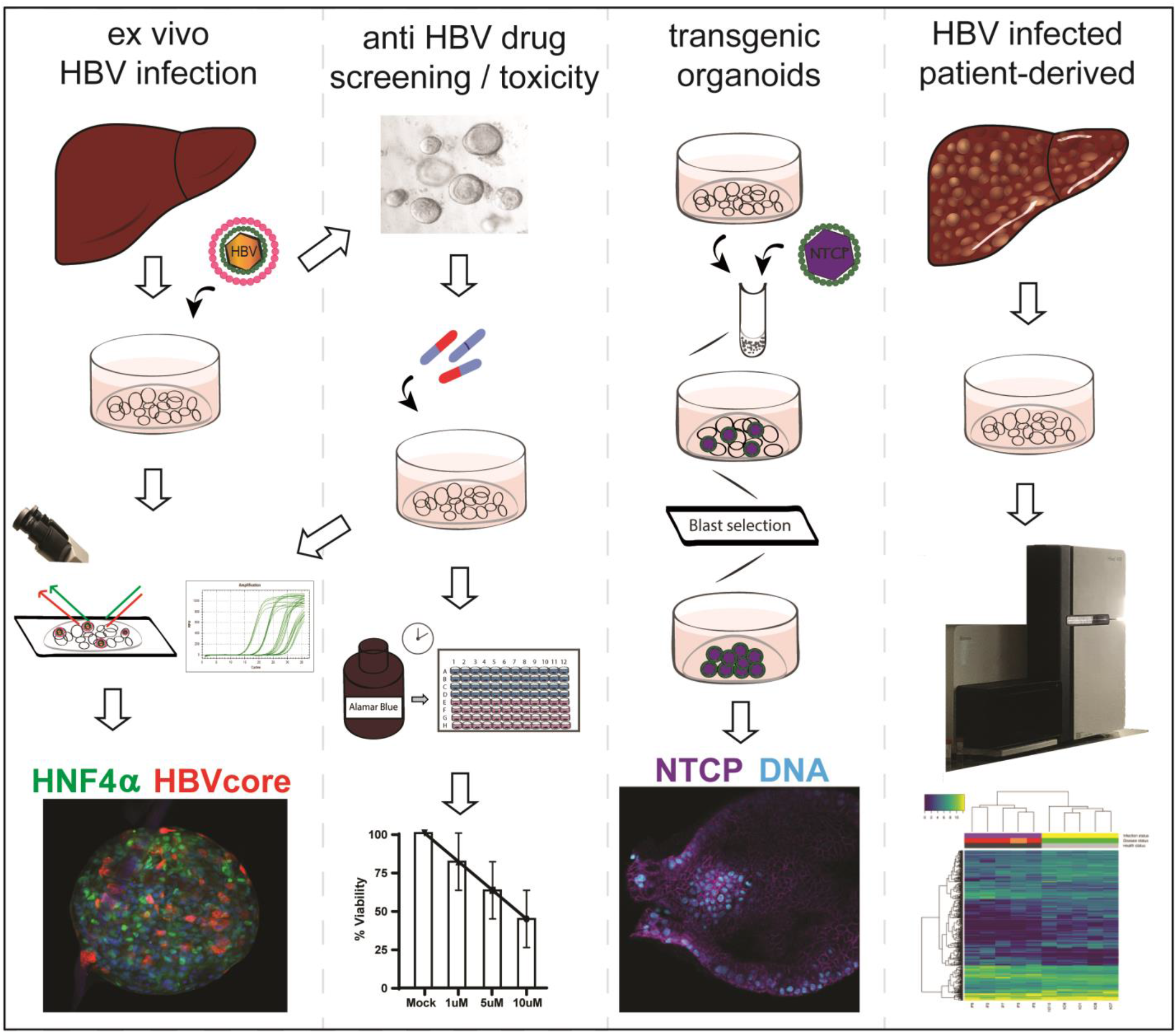
Schematic of applications of human liver organoids in HBV studies.

## Discussion

Human liver organoids are a novel in vitro primary model system that support HBV infection and replication, can be utilized as a platform to study HBV pathogenesis as well as for anti-HBV-drug screening, can be transgenically modified for downstream mechanistic and functional assays, and can be seeded from HBV-infected patient liver for personalized molecular and functional analysis (Figure 5). Healthy donor liver organoids were efficiently infected with both recombinant virus as well as HBV infected patient serum, expressed HBS Ag and HBV core proteins, produced cccDNA and infectious HBV in the culture supernatant indicting that liver organoids support the full HBV replication cycle (Figure 1). Human liver organoids therefore serve as an invaluable tool in the field of HBV-research to investigate the molecular mechanism underlying HBV replication in a primary cell system that is expandable and biobankable.

We demonstrated that the ex vivo infected organoid replication platform is amenable to screening for potential inhibitors of HBV replication and can provide insights into both HBV-directed anti-viral activity as well as drug-induced toxicity. The ability to measure different readouts of HBV replication at the level of HBV mRNA gene expression, or viral production by quantitation of HBV DNA or S Antigen present in the culture supernatant of HBV-infected liver organoids, can provide insight into the distinct steps in the HBV life cycle that may be targeted upon drug treatment. This is highlighted by our studies using the HBV-directed drugs Tenofovir and Fialuridine (Figure 2a), which as expected did not affect the levels of intracellular HBV RNA, but blocked HBV replication by inhibiting the reverse transcription step which converts HBV pre-genomic RNA to DNA, as we observed by decreased HBV DNA levels. A critical aspect of potentially therapeutic novel drugs which causes failure within the drug development pipeline is toxicity. While Fialuridine treatment of HBV infected human liver organoids efficiently inhibited HBV replication, it also caused significant toxicity in the liver organoids. The importance of this pre-clinical human liver organoid model to measure drug-induced toxicity is highlighted by the results of a phase II clinical trial conducted in 1993 to evaluate Fialuridine as a novel HBV-anti-viral drug (Eli Lilly Trial H3X-MC-PPPC, NIH protocol #93-DK-0031). Although the preliminary results of the trial were promising with reduced levels of HBV DNA in patients, the study was terminated on an emergency basis by week 13 because of serious hepatotoxicity in seven patients of which 5 died and two required emergency liver transplantation (McKenzie, 1995). This compound successfully passed pre-clinical toxicity studies, but the expression of a nucleoside transporter in human mitochondria may be responsible for the human-specific mitochondrial toxicity caused by Fialuridine (Lee, Lai, Zhang, & Unadkat, 2006). In the primary human liver organoid system, the toxicity of Fialuridine is significant and quantifiable (Figure 2e-f) but was not observed in the human HepG2 hepatoma cell line indicating that human liver organoids can more accurately predict clinical outcomes of drug treatment. Thus the HBV infected organoid platform can be used as an important pre-clinical, primary hepatocyte screening platform for both discovery of candidate new drugs targeting HBV transcription and replication, as well as their potential toxicity.

When cultured under differentiating growth conditions, the expression of the HBV receptor NTCP increased significantly in liver organoids. Consistent with the observed higher NTCP levels, in vitro infection of differentiated organoids with HBV led to higher levels of infection and replication. To obtain an expanding long term HBV infected replicating cultures of human liver organoids, using transgenic modification by lentiviral transduction, we generated liver organoids that exogenously expressed membrane-localized NTCP under expansion conditions. Surprisingly, the presence of NTCP did not result in more efficient infection of organoids, suggesting that although necessary, NTCP alone may not be sufficient for optimal infection (Figure 3). Consistent with our observations, HepG2 cell lines exogenously expressing NTCP, while susceptible to infection are inefficiently infected with HBV and require high viral titers (Iwamoto et al., 2014; Yan et al., 2012). These observations are suggestive of the potential necessity for additional (co)receptors or downstream factors necessary for optimal infection in hepatocytes. Using lentiviral transduction, we also generated polyclonal liver organoids that contain integrated full length HBV genome, which efficiently produced HBV under expansion conditions. This model, similar to HepG2.2.15 cell, although not a cccDNA generating model of productive HBV infection, is a useful primary human liver model to investigate HBV transcription and HBV-directed drugs.

Recently alternative liver organoid model systems derived from human (induced) pluripotent stem cells (Kaneko et al., 2016; Nie et al., 2018; Sakurai et al., 2017; Xia et al., 2017) as well as primary human hepatocyte coculture systems (March et al., 2015; Xiang et al., 2019)(Xiang et al., 2019) have been developed that can be used to study hepatitis B virus infection. However, the human liver organoid platform offers two main advantages over these model systems. The first and most important is that HBV infected patient-derived organoids resemble the diseased tissue of origin, as demonstrated by our gene expression studies (Figure 5). Second, the organoid platform allows the possibility of expansion and biobanking, ensuring the availability of sufficient samples for downstream analyses. Therefore, while primary human hepatocytes and iPSC models, similar to the ex vivo infection of hD human liver organoids we describe here, provide reliable ex vivo infection models for studies on HBV, the human liver organoid platform is uniquely suitable for long term, patient-derived studies.

Early surveillance of the changes in gene expression or biomarkers that predict the occurrence of HCC is critical to enable patients to receive timely and successful treatment. Paving the way for personalized therapy of HBV infected patients, liver organoids can be seeded from infected patient liver resection, at different stages of the disease. Comparison of the gene expression profiles of healthy donor to HBV-infected patient-derived liver organoids, and further HDV co-infected and acute liver failure patient sub-groups point to the extent to which organoids reflect the biology and represent the transcriptomic profiles of the primary non tumor tissue of origin. Interestingly, consistent with the absence of infection of liver stem cells from which organoid cultures are seeded, we found no sign of viral production in the non-tumor HBV infected patient-derived liver organoid lines. Thus, the observed HBV infected patient-specific gene expression profile is likely generated from chronic uninfected bystander cells. Our observations are consistent with the notion that the gene expression profile of the tumor microenviroment may serve as an important biomarker of HCC as bystander cells are chronically exposed to released HBV proteins despite their uninfected status. Indeed, our data indicates that HBV infected non tumor patient-derived liver organoids exhibit an early cancer gene profile, as shown by comparison to the TCGA-LIHC database. The identification of early aberrant gene regulatory networks and biomarkers that drive HBV-mediated HCC, detectable at an early non-tumor stage without phenotypic signs of tumorigenesis, allows for patient-specific surveillance of disease progression and provides a screening platform for candidate drugs that guide personalized treatment by targeting early stages in HBV-mediated HCC.

## Materials and Methods

### Liver tissue

The Medical Ethical Council of the Erasmus Medical Center approved the use of this material for research purposes, and informed consent was provided from all patients. Biopsies from twelve explanted HBV infected livers and from one HBV infected donor liver were fixed for 24 hours at room temperature in 4% formaldehyde solution (Klinipath) immediately after collection in the operating room. Fixed biopsies were processed according to standard protocol to dehydrate and infiltrate with paraffin wax and subsequent embedding into paraffin blocks. Sections of 4μm were cut using a microtome and mounted on glass microscope slides. After deparaffination according to standard procedure, the tissue slides were stained with haematoxylin and eosin (Haemalum – Mayer’s, VWR) according to the manufacturers protocol, dehydrated and mounted for microscopic analysis.

### Isolation and culture of human liver organoids

Liver organoids from healthy donors and HBV infected patients were isolated and cultured using the method previously described by Huch et al.(Huch et al., 2015) with minor modifications. In brief, liver specimens (1-2 cm^3^) were washed once with DMEM (Sigma) supplemented with 1% FCS and with 0.1% Penicillin Streptomycin (PS, Sigma), minced and incubated at 37°C with the digestion solution (collagenase 2.5mg/ml in EBSS). Incubation was performed for 30 minutes, further mincing and mixing the tissue every 10 minutes. To recover the cells, digestion solution was passed through a 70μM strainer in a 50 ml tube (GreinerBio) and washed with 45ml Advanced DMEM (Gibco) supplemented with 1% PS, 10mM HEPES (Gibco) and 1% GlutaMax (Gibco), henceforth Ad+++. Partially digested tissue was recovered from the strainer and further incubated with TrypLE Express (Thermoscientific) for 15 minutes at 37°C. Cells obtained from the first and second digestion were pooled together and washed twice with Ad+++. After the second centrifugation (200g, 5 minutes) cells were counted, mixed with an appropriate amount of BME solution (2/3 Basement Membrane Extract, Type 2 (Pathclear) diluted with 1/3 Ad+++) and seeded in 25μl drops containing 10000-15000 cells in 48 well suspension plates (GreinerBio). After BME solution had solidified, wells were filled with 250μl of human liver organoid isolation medium consisting Ad+++ supplemented of 1X B27 supplement without retinoic acid (Gibco), 1X N2 supplement (Gibco), 1.25mM N-acetyl-L-cysteine (Sigma), 20% (vol/vol) Rspo-1 conditioned medium(Huch et al., 2013), 1.25% (vol/vol) Wnt3a-conditioned medium(Barker et al., 2010), 10mM nicotinamide (Sigma), 10nM recombinant human (Leu15)-gastrin I (Sigma), 50ng/ml recombinant human EGF (Peprotech), 100ng/ml recombinant human FGF10 (Peprotech), 25ng/ml recombinant human HGF (Peprotech), 10μM Forskolin (Sigma), 5μM A8301 (Tocris), 25ng/ml Noggin (Peprotech) and 10μM Y27632 Rho Kinase (ROCK) Inhibitor (Sigma). After 1 week, isolation media was changed to human liver expansion media (EM; Ad+++ supplemented of 1X B27 supplement without retinoic acid (Gibco), 1X N2 supplement (Gibco), 1.25mM N-acetyl-L-cysteine (Sigma), 20% (vol/vol) Rspo-1 conditioned medium, 1.25% (vol/vol) Wnt3a-conditioned medium,(Barker et al., 2010) 10mM nicotinamide (Sigma), 10nM recombinant human (Leu15)-gastrin I (Sigma), 50ng/ml recombinant human EGF (Peprotech), 100ng/ml recombinant human FGF10 (Peprotech), 25ng/ml recombinant human HGF (Peprotech), 10μM Forskolin (Sigma) and 5μM A8301 (Tocris)) (Huch et al., 2015).

EM was changed twice a week, and cultures were split every 7-10 days according to organoid density. For passaging (1:4-1:8, depending on growth rate of the culture), organoids were resuspended in 10ml Ad+++, incubated in ice for 10 minutes and collected by centrifugation (5 minutes at 200g). Subsequently, organoids were incubated for 1-2 minutes in TrypLE Express at room temperature and mechanically disrupted by pipetting. After a further wash in Ad+++, cells were resuspended in BME solution and seeded in 24 or 48 wells suspension plates. After BME solution had solidified, wells were filled with 500μl (24 wells) or 250μl (48 wells) of human liver organoid expansion medium.

### Hepatic differentiation of liver organoids

Human liver organoid cultures derived from healthy and HBV infected livers were seeded and cultured for 4 days in EM without Wnt3a-conditioned medium supplemented with 25ng/ml of BMP7 (Peprotech). Hepatic differentiation was induced by culturing human liver organoids in differentiation medium (DM; Ad+++ supplemented with 1X B27 supplement without retinoic acid, 1X N2 supplement, 1mM N-acetylcysteine, 10nM recombinant human [Leu15]-gastrin I, 50ng/ml recombinant human EGF, 25ng/ml recombinant human HGF, 0.5μM A83-01, 10μM DAPT (Sigma), 3μM dexamethasone (Sigma), 25ng/ml BMP7 and 100ng/ml recombinant human FGF19 (Peprotech)). Differentiation medium was changed twice a week for 7 days before infection or 10 days before staining for Albumin and HNF4α(Huch et al., 2015). For subsequent downstream analysis the following amounts of organoids are required (Table 4).

**Table 4.** Recommended starting material quantities.

| Procedure | Number of wells/ condition | Number of organoids/ well |
| --- | --- | --- |
| DNA isolation | 1x 24 well plate | ≈100-500 organoids/well |
| RNA isolation | 1x 24 well plate |  |
| cccDNA isolation | 20-24x 24 well plate |  |
| Immunofluorescence staining | 5-10x 24 well plate |  |

### Total RNA isolation and quantitative RT qPCR

RNA extraction was performed starting from 1-2 wells of a 24 wells plate, organoids were collected in 500μl of cell lysis buffer and processed using RealiaPrep RNA Cell Miniprep System (Promega), according to manufacturer’s instructions. cDNA synthesis was performed starting from 300-1000ng of RNA using Superscript II Reverse Transcriptase (Life Technologies) kit following manufacturer’s protocol for random primer cDNA synthesis. cDNA was diluted 1:5 in nuclease free water and 2μl of the diluted product was used for real time PCR with the following reagents: 5μl of GoTaq qPCR Master Mix (Promega), 2μl of nuclease free water and 1μl of 10mM primer mix. Amplification was performed on the CFX Connect Real-Time PCR Detection System thermocycler (BioRad) using following thermal program starting with 3 minutes at 95°C, followed by 40 cycles of 95°C for 10 seconds and 60°C for 30 seconds. Specificity of the RT-qPCR products was assessed by melting curve analysis. Primers used for real-time qPCR are listed below.

GAPDH Fwd 5’-CAAGAAGGTGGTGAAGCAG-3’; Rev 5’-GCCAAATTCGTTGTCATACC-3’
KRT7 Fwd 5’-CTCCGGAATACCCGGAATGAG-3’; Rev 5’-ATCACAGAGATATTCACGGCTCC-3’
CYP3A4 Fwd 5’-TGTGCCTGAGAACACCAGAG-3’; Rev 5’-GTGGTGGAAATAGTCCCGTG-3’
NTCP Fwd 5’-GCTTTCTGCTGGGTTATGTTCTC-3’; Rev 5’-CATCCAGTCTCCATGCTGACA-3’
HNF4A Fwd 5’-CGTGCTGCTCCTAGGCAATGAC-3’; Rev 5’-ACGGACCTCCCAGCAGCATCT-3’
SOX9 Fwd 5’-GGAAGTCGGTGAAGAACGGG-3’; Rev 5’-TGTTGGAGATGACGTCGCTG-3’
ALBUMIN Fwd 5’-GCGACCATGCTTTTCAGCTC-3’; Rev 5’-GTTGCCTTGGGCTTGTGTTT-3’
LGR5 Fwd 5’-AGGTCTGGTGTGTTGCTGAGG-3’; Rev 5’-TGAAGACGCTGAGGTTGGAAGG-3’
LDLR Fwd 5’-GACGTGGCGTGAACATCTG-3’; Rev 5’-CTGGCAGGCAATGCTTTGG-3’
PCSK9 Fwd 5’-AGGGGAGGACATCATTGGTG-3’; Rev 5’-CAGGTTGGGGGTCAGTACC-3’

Gene expression levels were calculated using the 2ΔCt method, whereas fold increase was calculated using the 2ΔΔCt (Schmittgen & Livak, 2008). GAPDH was used as housekeeping control.

### Immunofluorescence and image analysis

Human liver organoids were collected and washed three times to with cold Ad+++ to remove BME, then fixed with 4% paraformaldehyde for 30 minutes in ice and permeabilized using 0.3% (HBcAg, HNF4α and NTCP) or 1% (Albumin) Triton X-100 (Sigma) in PBS for 30 minutes at room temperature. For HBsAg staining cells were fixed and permeabilized in 100% Acetone. Specimens were incubated for 2h at room temperature in PBS plus 10% BSA (Roche) and 0.5% FCS (HBsAg) or PBS plus 0.5% FCS, 0.3% triton, 1% BSA 1%DMSO (Albumin, HBcAg, HNF4α and SLC10A1). Following blocking, human liver organoids were incubated overnight with primary antibodies (mouse anti-HBcAg, mouse anti-HBsAg (ThermoScientific), goat anti-Albumin, rabbit anti-HNF4α (Santa Cruz Biotechnology) and rabbit anti-SLC10A1 (NTCP, Sigma) diluted in PBS + 10% blocking buffer. After extensive washing, human liver organoids were stained with appropriate Alexa Fluor dye-conjugated secondary antibodies (Life Technologies). Nuclei were stained with Hoechst33342 (Molecular Probes). Immunofluorescence images were acquired using a confocal microscope (Leica, SP5). Images were analyzed and processed using Leica LAS AF Lite software (Leica SP5 confocal). All phase contrast pictures were acquired using a Leica DMIL microscope and a DFC420C camera.

### Production of HBV virus and HBV infection

HepG2.2.15 cells, a HepG2 derived cells line stably transfected with full length HBV (kindly provided by Prof. Bart Haagmans, Erasmus MC) were cultured in DMEM medium (Gibco) supplemented with 10% fetal bovine serum (Gibco) and 1% penicillin/streptomycin. For virus production, 3×10^6^ cells were plated in collagen coated 10cm plates, cultured in supplemented DMEM until confluency and subsequently in Ad+++ for 4 days. The supernatant of HepG2.2.15 cells was then collected, filtered and concentrated using the PEG Virus Precipitation Kit (Abcam) following the manufacturer’s instructions. Precipitated virus was aliquoted and stored at −80°C until use. Human serum was obtained from residual samples from HBV infected individuals attending Erasmus MC for routine clinical activity. As negative control, an aliquot of the virus equivalent to the inoculum was inactivated by incubation at 100°C for 30 minutes. Human liver organoids were resuspended using either active virus or heat inactivated control at an MOI of 1-10 * 10^3^ copies HBV DNA/organoid, transferred to 24 wells plate and centrifuged for 1h at 600g. Following spinoculation, plates were incubated at 37°C for 5h and then seeded in BME following the culturing protocol. After BME solution has solidified, liver organoids were maintained in EM for 16h, washed 4 times with Ad+++ and cultured EM or DM as indicated.

### Detection of HBV DNA and RNA

DNA was extracted from culture supernatants using the QIAamp MinElute Virus Spin Kit following manufacturer’s instructions. DNA extracted from supernatant and cDNA obtained from reverse transcription of intracellular DNA (see section 9 for details) were analyzed in duplicate using a TaqMan based qPCR assay. For each reaction, a 25ul mixture was prepared containing 2.5μl 10X Buffer, 1.75μl 25mM MgCl2, 1μl 10mM dNTPs, 1U of Platinum Taq, 0.125μl of 100μM forward (5’-GCAACTTTTTCACCTCTGCCT A-3’) and reverse primer (5’-AGTAACTCCACAGTAGCTCCAAATT-3’), 0.075μl of 50μM probe (FAM-TTCAAGCCTCCAAGCTGTGCCTTGGGTGGC-TAMRA), and 7.5μl (DNA) or 4μl (cDNA) of template. Each PCR reaction included a standard curve made of dilutions of a plasmid containing the full-length HBV genome ranging from 4 to 4×10^5^ copies of plasmid. Beta-2-microglobulin was used as housekeeping control for expression analysis of cDNA samples (B2M Fwd 5’-AGCGTACTCCAAAGATTCAGGTT-3’, B2M Rev 5’-ATGATGCTGCTTACATGTCTCGAT-3’, B2M probe FAM-TCCATCCGACATTGAAGTTGACTTACTG-BHQ1).

Detection of pol and HBX transcripts was performed using a previously published nested PCR protocol (Wong et al., 2011).

### Isolation and detection of HBV cccDNA from infected liver organoids

HBV cccDNA was isolated from HBV and HI (negative control) infected human liver organoids by a previously described alkali lysis plasmid DNA isolation protocol with a minor modification (Yang, Mason, & Summers, 1996). Briefly, Human liver organoids were collected on day 8 post infection incubated on ice for 10-20 mins and washed with Ad+++ to remove the BME. After centrifugation (5 mins at 1000rpm) organoids were treated with TrypLE Express and incubated for 30-50 sec at room temperature. Organoids were washed with PBS and collected by centrifugation (5 mins at 1000rpm), resuspended with 800μl of ice-cold cell lysis buffer (1mM EDTA (pH 8.0), 5mM Tris: HCl (pH 7.5) and 0.05% Nonidet P-40). After 10 min incubation on ice, an equal volume of alkali lysis buffer (0.1M NaOH, 6% SDS) was added and the solution was incubated for 30 mins at 37°C. DNA was neutralized by adding 3M potassium acetate (pH 5.0) to a final concentration of 0.6M and centrifuged for 5 mins at 12000 rpm. The supernatant was extracted two times with phenol followed by extraction with butanol: isopropanol (7:3) for removal of residual phenol. Subsequently, the DNA was precipitated with 1ml 100% ethanol, 400μl 7.5M ammonium acetate and 1μl 20mg/ml glycogen overnight at −80°C. Next day, the cccDNA sample was spun down for 30 mins at 4°C, 14000 rpm and washed with 70% ethanol. After spinning the samples for 15 mins at 4°C, 14000 rpm, the pellet was resuspended in 50μl of nuclease free water.

To remove the chromosomal DNA or any linear HBV DNA, 25μl of the isolated HBV cccDNA, HBV plasmid (positive control) and HBV DNA from inoculum was digested with plasmid-safe DNase (Epicentre, E3101K, Madison, WI). According to the manufacturer’s protocol, the samples were digested for 1 hr at 37°C followed by 30 mins of heat inactivation at 70°C. The digested samples were purified once with phenol: chloroform: isoamyalcohol (Sigma-Aldrich) followed by treatment chloroform: isoamylalcohol (24:1), cccDNA was precipitated with 20ul 3M NaAC (pH 5.2), 1ul glycogen (20mg/ml), 1ml 100% ethanol and washed as described above.

For quantification of cccDNA a TaqMan based qPCR was performed, whereby for each reaction 20μl reaction-mix was prepared containing 4.2μl of the diluted cccDNA, 10μl LightCycler®480 Probes Master (Roche), 1μM primer mix (Fwd 5’-GTCTGTGCCTTCTCATCTGC-3’; Rev 5’-AGTAACTCCACAGTAGCTCCAAATT-3’), 0.2μM probe (FAM-TTCAAGCCTCCAAGCTGTGCCTTGGGTGGC-BHQ1) and 4% DMSO. qPCR was carried out using a previously published protocol 95°C for 10 min, followed by 50 cycles of 95°C for 15 s, and 61°C for 1 min (Winer et al., 2017).

### Infection with HBV generated from organoids

the supernatant produced by infected organoids at 4-8 days post infection was collected and concentrated using Amicon Ultra-15 100K (Milipore). Human liver organoids were resuspended using either concentrated HBV virus or HI control. After an hour of spinoculation at 32°C 600g, plate was incubated at 37°C overnight and then washed & seeded in BME following the HBV infection and culturing protocol.

### Cell culture

HepG2 and HepG2.215 cells were cultured in DMEM medium (Gibco) supplemented with 10% fetal bovine serum (Gibco) and 1% penicillin/streptomycin and incubated in 5% CO2 at 37°C.

### Viability assays

HepG2 cells with cell density of 2×10^4^ cells/ml were seeded in 24-well plate with DMEM high-glucose media supplemented with fetal bovine serum (10% v/v) and penicillin/streptomycin (1% v/v) and incubated in 5% CO2 at 37 °C for overnight. Human liver organoids seeded from healthy donor or HBV infected patient livers were split in the ratio of 1:10, seeded in 20μl BME 3D culture in 48- well plates and differentiated as described above. 10 days post differentiation for human liver organoids and 24 hrs post seeding of the HepG2 cells they were treated with different concentrations of Fialuridine (Cayman, 15867-1) or Tenofovir disproxil fumarate (Sigma, SML1794) (1-20μM) or the vehicle control. At different time points (2-10 days) post treatment, organoid viability was measured using the AlamarBlue viability assay (alamarBlue™ Invitrogen DAL1025, 1:10 in DM) according to manufacturer’s instructions. Briefly, treatment DM medium was removed from wells and 10% Alamar blue with DM was added to each well and incubated for 4 hours (organoids) or 2 hours (HepG2 cells) at 37°C before absorbance readings were taken at 570 and 600 nm. The results were normalized to control vehicle treated differentiated organoids. The lower limit of quantitation of the assay was determined by values obtained from BME without organoids and is represented by dotted line in figure panels. Each treatment condition was repeated at least 3 times and readings were done in duplicate. Cell imaging was performed after each Alamar blue assay.

### 3’ mRNA sequencing

For each RNA preparation, 2 wells of organoids in expansion phase were collected in Trizol reagent (Sigma) 4-5 days after splitting. RNA was extracted according to manufacturer’s instruction and resuspended in 30μl of nuclease free water. Total RNA was quantitated (ND1000 Spectrophotometer – PEQLAB). Samples were diluted accordingly to a mean concentration of approximately 100-150ng/μl and their quality assessed on a Bioanalyzer (Agilent Technologies) using the Agilent RNA 6000 Nano Kit reagents and protocol (Agilent Technologies). RNA samples were processed for library preparation using the 3’ mRNA-Seq Library Prep Kit Protocol for Ion Torrent (QuantSeq-LEXOGEN™, Vienna, Austria), according to manufacturer’s instructions. Briefly, up to 500ng from each RNA sample were used for first strand synthesis. The RNA was subsequently removed and 2nd strand synthesis was initiated by a random primer, containing Ion Torrent compatible linker sequences and appropriate in-line barcodes. 2nd strand synthesis was followed by magnetic bead-based purification and the resulting library was PCR-amplified for 14 cycles and re-purified. Library quality and quantity was assessed on a Bioanalyzer using the DNA High Sensitivity Kit reagents and protocol (Agilent Technologies). The quantified libraries were pooled together at a final concentration of 100pM. The pools were templated and enriched on an Ion Proton One Touch system. Templating was performed using the Ion PI™ Hi-Q™ OT2 200 Kit (Thermo Fisher Scientific), followed by sequencing using the Ion PI™ Hi-Q™ Sequencing 200 Kit on Ion Proton PI™ V2 chips (Thermo Fisher Scientific), an Ion Proton™ System, according to the manufacturer’s instructions.

### Short read mapping

The Quant-Seq FASTQ files obtained from Ion Proton sequencing were mapped on the UCSC hg19 reference genome using a two-phase mapping procedure. Firstly, the short reads were mapped using tophat2(Kim et al., 2013), with the following non-default settings: --read-mismatches 3 --read-gap-length 3 --read-edit-dist 3 --no-novel-juncs, other settings at default. Additional transcript annotation data for the hg19 genome from Illumina iGenomes (http://cufflinks.cbcb.umd.edu/igenomes.html) was also provided to tophat2 for guidance. Next, the reads which remained unmapped were converted back to FASTQ files using bam2fastq from the BEDTools (Quinlan & Hall, 2010) suite and submitted to a second round of mapping using Bowtie2 (Langmead & Salzberg, 2012) against the hg19 genome with the --local and --very-sensitive-local switches turned on. All resulting BAM files were visualized in the UCSC Genome Browser using BEDTools and tools provided by the UCSC Genome Browser toolkit.

### Statistical analysis of Quant-Seq data

The resulting Quant-Seq BAM files were analyzed with the Bioconductor package metaseqR(Moulos & Hatzis, 2015) which has built-in support for Quant-Seq data. Briefly, the raw BAM files, one for each organoid sample, were summarized to a 3’ UTR read counts table from Ensembl longest (dominant) transcripts (version 90). The original 3’ UTR regions were extended 500bp upstream and downstream to accommodate the variable read length of Ion Proton reads. In the resulting read counts table, each row represented one 3’ UTR region, each column one Quant-Seq sample and each cell the corresponding read counts associated with each row and column. The final 3’ UTR read counts table was normalized for inherent systematic or experimental biases using the Bioconductor package DESeq (Anders, Reyes, & Huber, 2012) after removing areas that had zero counts over all the Quant-Seq samples. Prior to the statistical testing procedure, the 3’ UTR read counts were filtered for possible artifacts that could affect the subsequent statistical testing procedures. 3’ UTR areas presenting any of the following were excluded from further analysis: i) 3’ UTR areas corresponding to genes smaller than 500bp, ii) 3’ UTRs with read counts below the median read counts of the total normalized count distribution. Similar expression thresholds (e.g. the median of the count distribution) have been previously used in the literature(Mokry et al., 2012), where the authors use the median RPKM value instead of normalized counts), iii) 3’ UTR areas corresponding to genes with the following Ensembl biotypes: rRNA, TR_V_pseudogene, TR_J_pseudogene, IG_C_pseudogene, IG_J_pseudogene, IG_V_pseudogene. The remaining 3’ UTR counts table after filter application was subjected to differential expression analysis for the appropriate contrasts using the PANDORA algorithm implemented in metaseqR. 3’ UTR areas (and their corresponding genes) presenting a PANDORA p-value less than 0.05 and fold change (for each contrast) greater than 1 or less than −1 in log2 scale were considered as differentially expressed.

### Clustering analysis

Hierarchical clustering was performed using the Euclidean distance and complete linkage for the construction of the dendrograms. The expression values used to generate the heatmap were DESeq-normalized read counts in log2 scale. Multidimensional scaling was performed using the Spearman correlation distance metric on the gene expression matrix using DESeq-normalized read counts in log2 scale. All calculations and visualizations were performed using facilities from the R language. All analysis scripts and logs of the analysis pipelines are available upon request.

### Gene Ontology and Pathway analysis

Gene Ontology (GO) enrichment and biochemical pathway analysis was performed using GeneCodis(Tabas-Madrid, Nogales-Cadenas, & Pascual-Montano, 2012). For the GeneCodis GO and pathway analysis the 361 iP “Signature” genes were used

### Generation of the lentiviral vectors and transduction of liver organoids

A gene block fragment encoding human NTCP was designed based on the reference sequence retrieved from NCBI nucleotide database (NM_003049.3). A 3X Flag PCR fragment including NTCP coding sequence was amplified using the NTCP_FWD (CACCATGGATTACAAGGATGACGACGATAAGGATTACAAGGATGACGACGATAAGGATTACAAGGATGACGACGATAAGATGGAGGCCCACAACGCGTCTgcccca) and NTCP_REV (TTACTAGGCTGTGCAAGGGGAGCA) primers and cloned in the pENTR/D-TOPO entry vector (Invitrogen) following manufacturer’s instructions.

A pENTR/D-TOPO vector harboring a competent full-length copy of HBV (1.3 wt genomes) was generated using a Gibson Assembly protocol. Two fragments encompassing 1.3 times the wt HBV genome were purified using the QIAquick PCR Purification Kit (Qiagen) after PCR using the following primer pairs: VectorFwd (5’-caaaaaagcaggctccgcggccgcccccttcacGGACGACCCTTCTCGGGG-3’)/MiddleRev (5’-gagaagtccaccacgAGTCTAGACTCTGCGGTATTGTGAG-3’) and MiddleFwd (5’-cgcagagtctagactCGTGGTGGACTTCTCTCAATTTTC-3’)/ VectorRev (5’-tgccaactttgtacaagaaagctgggtcggAGGGGCATTTGGTGGTCTATAAG-3’). The HepG2.2.15 DNA was used as template for the PCR: An empty pENTR/D-TOPO was cut with NotI and BssHI and gel purified using QIAquick Gel Extraction Kit (Qiagen). After purification, the entry vector and the PCR fragments were assembled using the Gibson Assembly Master Mix (New England BioLabs), following manufacturer’s instructions.

To generate lentiviral constructs, the fragments cloned into entry vectors were transferred to a lentiviral expression destination vector (pLenti6/V5-DEST Gateway Vector, Invitrogen), using the Gateway technology (Invitrogen). The full-length HBV pLenti6/V5-DEST vector was further modified in order to remove the CMV promoter. The vector was digested with two restriction enzymes flanking the CMV promoter, ClaI and PstI (New England Biolabs), and purified from gel using QIAquick Gel Extraction Kit. After filling end gaps using Klenow polymerase (New England Biolabs), plasmids were ligated and transformed in One Shot® Stbl3™ Chemically Competent E. coli (Invitrogen). All HBV and NTCP lentiviral vectors were sequenced to verify vector structure and integrity of open reading frames.

The lentiviral constructs were generated using the ViraPower Kit (Invitrogen) and 293FT cells, following manufacturer’s protocol. Briefly, one day prior transfection, 3×10^5^ cells were plated in 10cm dishes in DMEM + 10% FCS. The following day, DMEM was replaced with 5ml of Opti MEM I (Invitrogen) and cells were transfected with 9μg of the ViraPower Packaging Mix and 3μg of the pLenti6/V5-DEST Gateway Vector using Lipofectamine (Invitrogen). The day after transfection media was changed to DMEM + 10% FCS. Cell supernatant containing the lentiviral particles was collected 60 and 72 hours after transfection, filtered with a 0.42μm filter, aliquoted and stored at −80°C.

Early passage (passage 0 to 3) human liver organoids were collected in cold Ad+++ and incubate for 10 minutes in ice to remove BME. After centrifugation (5 minutes at 200g), organoids were resuspended in TrypLE Express and incubated until single cells were >80%. Cells were collected by centrifugation (5 minutes at 200g), resuspended in 1ml of lentiviral harvest and divided over 4 wells of a 48 wells plate. Plates were centrifugated for 1h at 600g and then incubated for 5h at 37°C. Afterwards, cells were collected by centrifugation (5 minutes at 200g), resuspended in BME solution and plated. After BME has solidified, wells were filled with EM supplemented with 25ng/ml Noggin and 10μM Y27632. Four days after infection medium was changed to EM supplemented with 25ng/ml Noggin and 10μM Rho Kinase (ROCK) Inhibitor Y27632 and 5ug/ml blasticidin. Organoids were kept under selection for 7 days and then media was changed to regular EM. Once selected organoids have recovered, cultures were splitted to remove dead cells and cultured according to the regular protocol. Parallel infections were performed using HepG2 cells and the same lentiviral constructs in order to generate control cell lines.

To determine NTCP functionality following transduction with NTCP lentiviral vector, parental and selected NTCP transduced organoids were cultured in EM or EM supplemented with different concentrations of Aruvastatin and Rosuvastatin for 12h. Organoids from 2 wells of a 24 wells plate were collected in 500μl of cell lysis buffer and processed using RealiaPrep RNA Cell Miniprep System (Promega), according to manufacturer’s instructions.

To determine HBV production following transduction with full length HBV lentiviral vector, supernatant and cells from transduced cultures were collected at different time points after completion of blasticidin selection. Presence of viral DNA in the supernatant and cellular associated viral RNA was assessed using the real time protocol detailed in section 11. Presence of HBsAg in organoid supernatant was assessed using the MonaLisa Kit (Promega) according to manufacturer’s instructions.

## Data availability

Sequencing data that support the findings of this study have been deposited in GEO with the accession code GSE 126798.

## Statistical analysis

The data were first analyzed by ANOVA, and then each pair was compared through Dunnett’s multiple comparisons test. A value of p < 0.05 was considered statistically significant. Data are shown as mean ± SD of at least 3 replicate treatments \**P* < 0.05; \*\**P* < 0.01. The data were analyzed and graphs were depicted by GraphPad Prism software 5 (GraphPad Software, La Jolla, CA, USA). Obtained data from drug screening of infected organoid have 2-8 technical replicates for each donor. Thus, the obtained data from each donor were analyzed by ANOVA separately.

## Acknowledgments

We would like to thank Vaggelis Harokopos for NGS and Karien Hamer for technical support.

TM received funding from the European Research Council (ERC) under the European Union’s Seventh Framework Programme (FP/2007-2013)/ERC STG 337116 Trxn-PURGE, Dutch AIDS Fonds grants 2014021 and 2016014, and Erasmus MC mRACE research grant.

## Author Contributions

EDC, SR, FC, MMK, AB, SGR, TWK, FP and TM carried out the experiments and performed data analysis. PM, CK, RJP, and PH performed RNAseq and gene expression analysis. MMA, LVDL, JMNI, CB provided tissue samples and serum from HBV infected patients. FP, MMA, LVDL, CB, HG, MH, SFB, SR, SGR, RV, HC, and provided expertise, material and contributed to the writing of the manuscript. EDC, FC, RV, PH and TM conceived the study and wrote the manuscript. All authors read and approved the final manuscript.

---

**Figure1-figure supplement 1:**
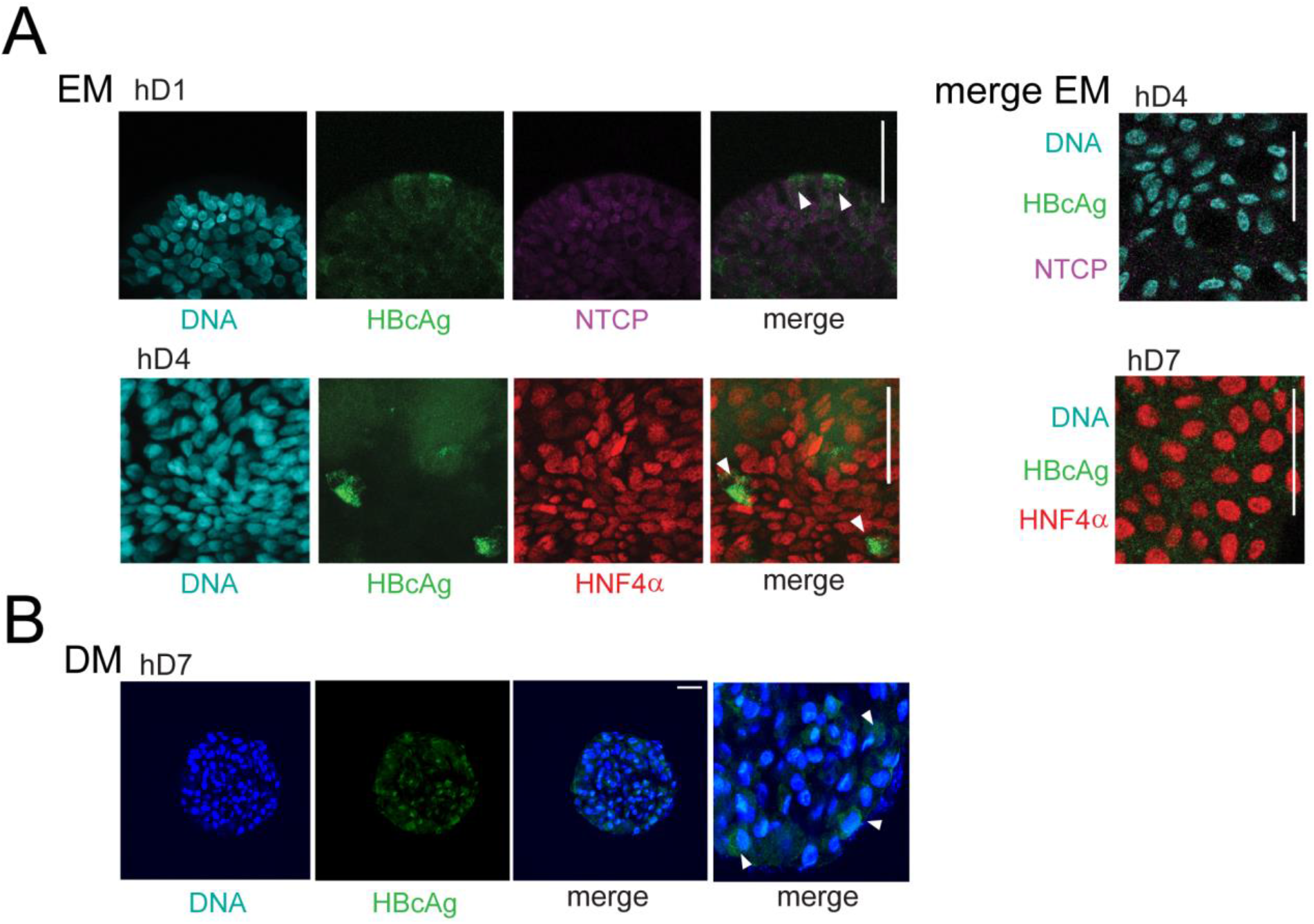
Expression of HBV core Ag in infected human liver organoids. (A) Immunofluorescent staining showing the expression of HBcAg (green) together with NTCP (magenta) or HNF4α (red) performed in different hD organoids 6 days after HBV infection in expansion condition (EM). (B) Representative immunofleorescent images of differentiated organoid (DM) showing the expression of HBcAg (green) and β-catenin (gray) 6 days after HBV infection.

**Figure 1 – figure supplement 2:**
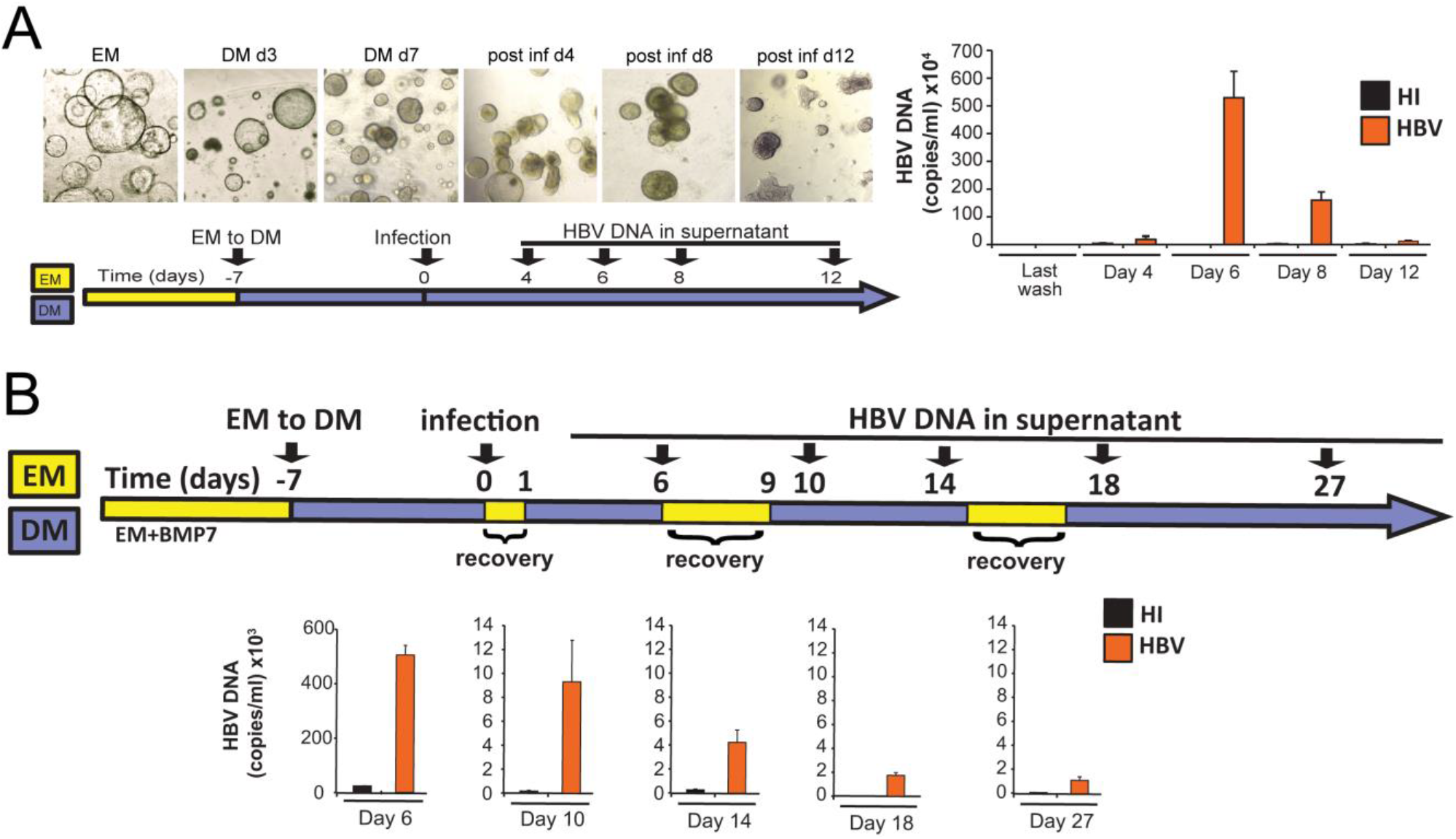
Release of HBV DNA declines over time in hD organoid lines infected in vitro. (A) Bright field images and levels of HBV DNA in the supernatant of infected organoids at different days post infection. (B) HBV DNA in the supernatant of infected organoids undergoing short EM pulse treatments after infection. HBV DNA was quantified by real time PCR and compared to the mock infected (heat inactivated, HI) cultures.

**Figure 1 – figure supplement 3:**
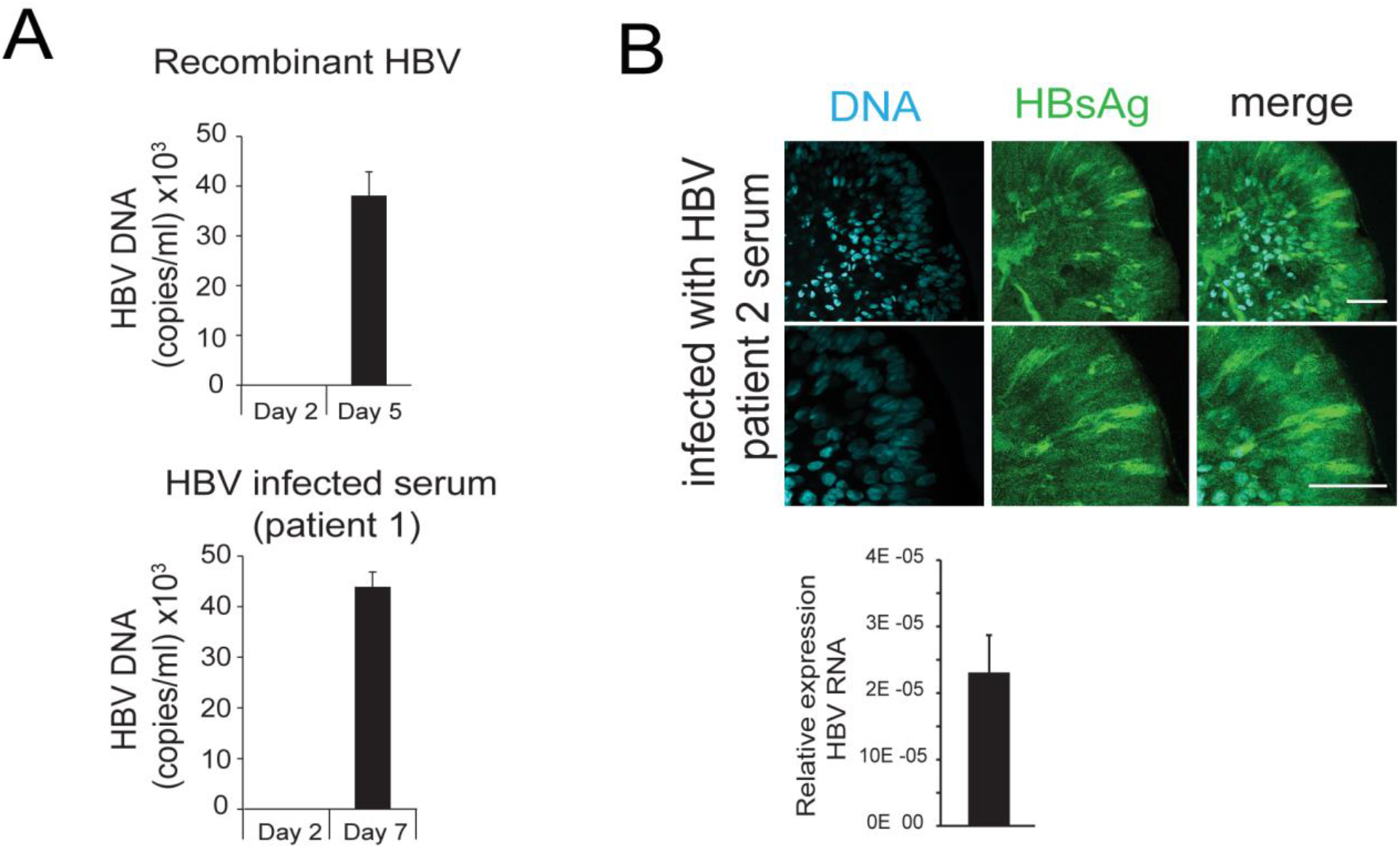
HBV specific DNA, RNA and proteins are detected in hD derived organoids upon infection with recombinant HBV virus as well as with HBV infected patient’s serum. (A) Levels of HBV DNA in supernatant of hD organoids infected with either recombinant virus derived from HepG2.2.15 cells or serum obtained from HBV positive individuals (B) Immunofluorescent staining showing the expression of HBV Surface Ag (green) in DM organoids 5 days after infection with serum obtained from HBV positive individuals (patient 2), scale bars represent 50 μm. Bar graph shows total HBV RNA levels in the culture at the time of staining.

**Figure 2 – figure supplement 1:**
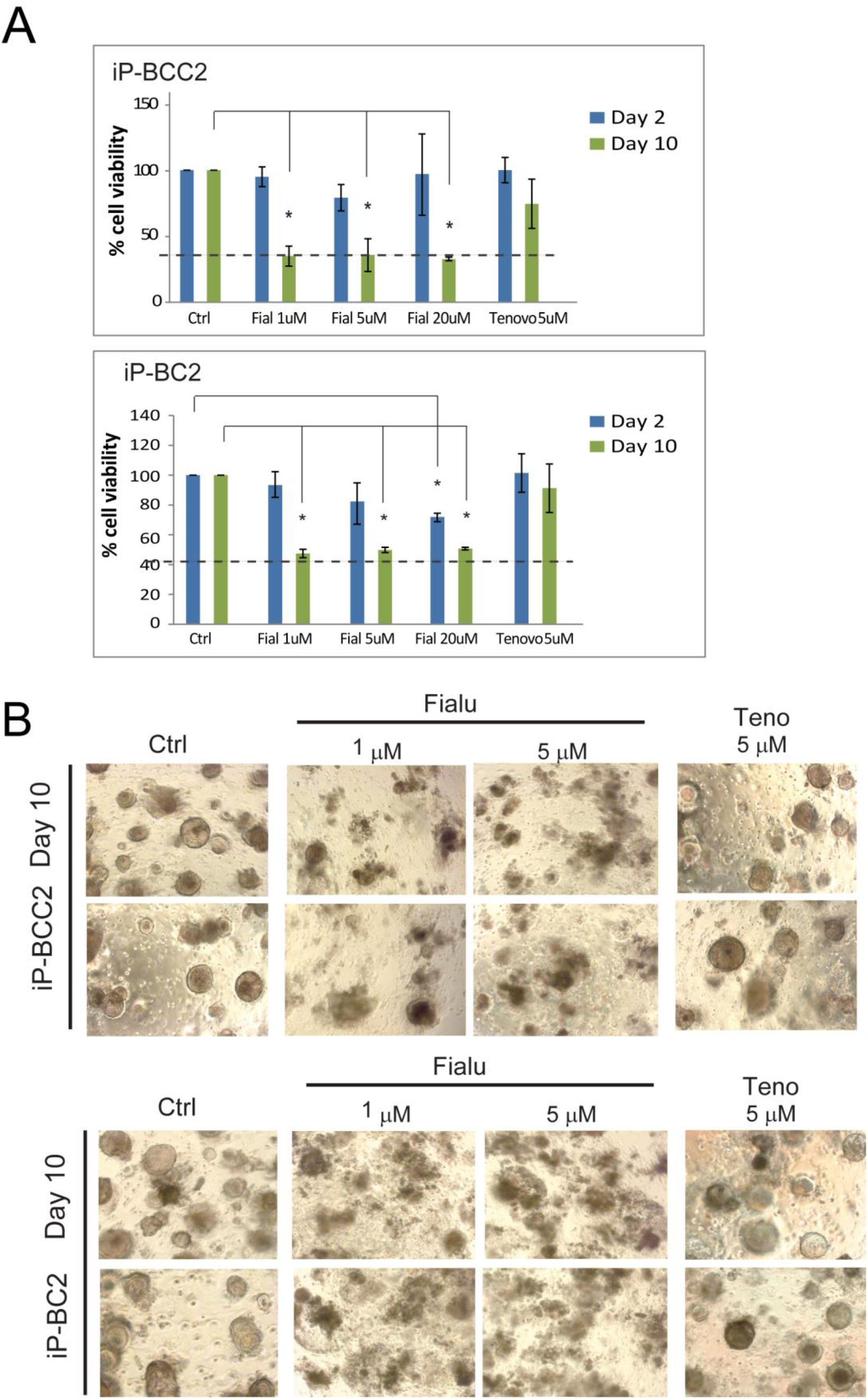
Human liver organoids as a model for anti-HBV drug-induced toxicity screening (A) Bar diagrams representing relative cellular viability of differentiated patient-derived liver organoids (iP-BCC2 and iP-BC2) after 2 or 10 days of treatment with Fialuridine or Tenofovir at the different concentrations indicated using the AlamarBlue cell viability assay. All values are normalized to the vehicle treated control and plotted as the average of percent viability ± SD (n=3). The dotted line represents the lower limit of quantification based on values obtained from wells free of organoids containing the BME matrix only. (B) Representative bright field images taken of differentiated patient-derived liver organoids (iP-BCC2 and iP-BC2) as indicated with Fialuridine and Tenofovir.

**Figure 3 – figure supplement 1:**
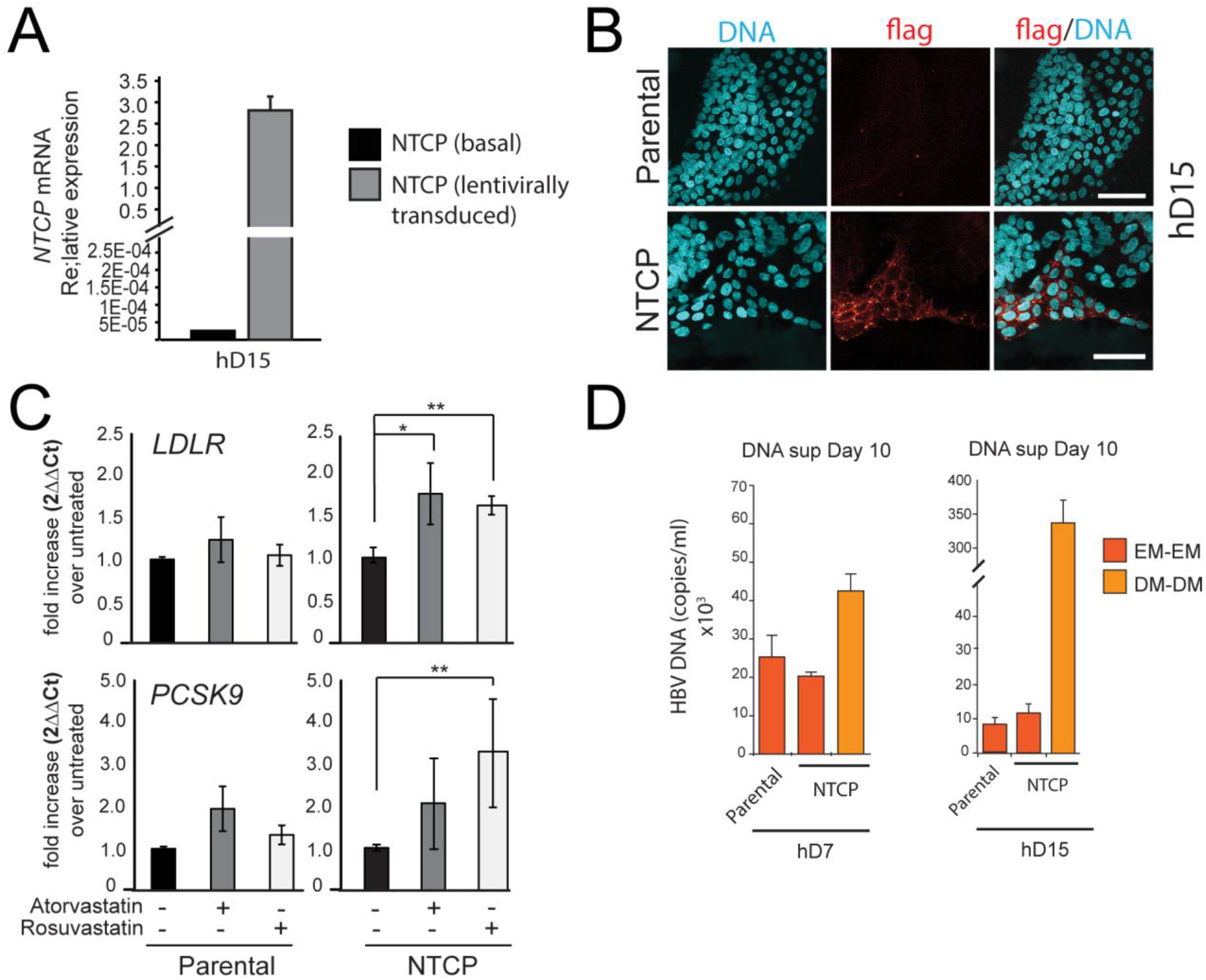
hD organoid lines expressing high levels of functional NTCP can be generated by lentiviral infection. (A) Levels of expression of NTCP were evaluated by RT in the untransduced (Parental) and the transduced (NTCP) lines. Expression of NTCP was calculated according to the 2ΔCt method using the housekeeping gene GAPDH as reference. (B) Expression of NTCP was confirmed by immunofluorescence staining using antibodies against Flag (red). (C) Evaluation of changes in cholesterol metabolism genes following transgenic expression of NTCP in liver organoids. The levels of LDLR and PCSK9 were evaluated by real time PCR following treatment with Atorvastatin and Rosuvastatin. Fold difference in gene expression was calculated according to the 2ΔΔCt method using untreated cells as reference. (D) HBV DNA levels in supernatant of parental cultures grown in EM and NTCP cultures (of donors hD7 and hD15) grown in EM or DM were quantified by real time PCR 10 days after infection.

**Figure 3 – figure supplement 2:**
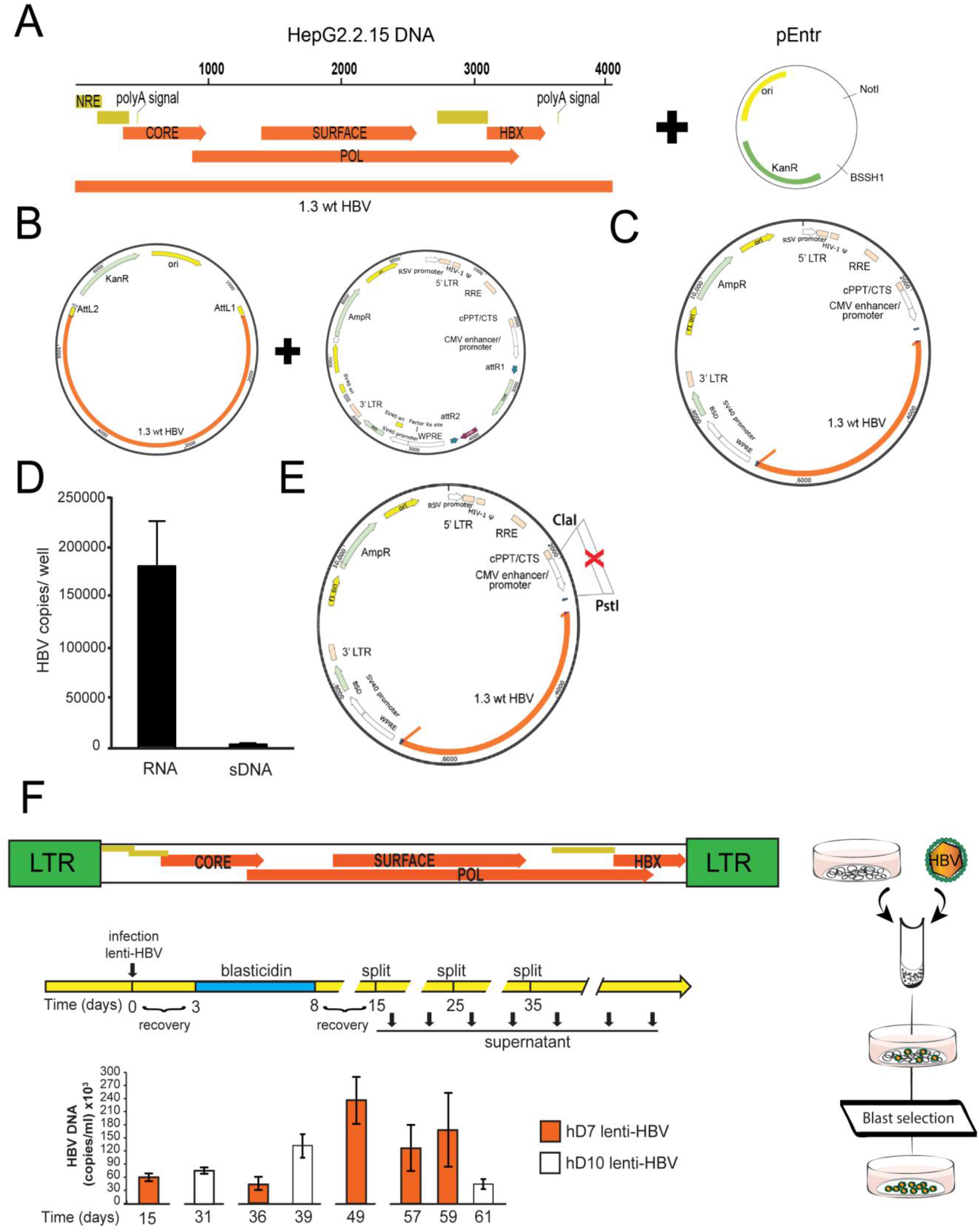
Generation of lentiviral construct used to establish human liver organoid lines expressing viral RNA and releasing virus in the supernatant. (A-B) A fragment encoding 1.3 times the wt HBV genome was amplified from HepG2.2.15 and cloned into the pEntr plasmid. (C) Using the gateway system, HBV was then transferred to a lentiviral expression vector under the control of CMV promoter. (D) Bar charts represent the amount of cellular associated HBV RNA and of HBV DNA released in the supernatant (sDNA) expressed as copies/well of culture. Although HBV RNA could be clearly detected after infection of organoids with the lentiviral construct generated in (B), viral release was minimal, likely because of transcriptional interference due to presence of the strong CMV promoter upstream of HBV genome, which was then removed from the construct (E). (F) Experimental procedure for the generation of transgenic organoid lines expressing full length HBV. EM organoids were infected with a lentiviral vector including a construct encoding 1.3 copies of the HBV genome and selected with blasticidin for 5 days in order to obtain stable transgenic organoid lines. Viral production was determined by real time PCR, measuring the amount of HBV DNA secreted in the supernatant at regular intervals and up to 62 days after lentiviral infection.

**Figure 4 – figure supplement 1:**
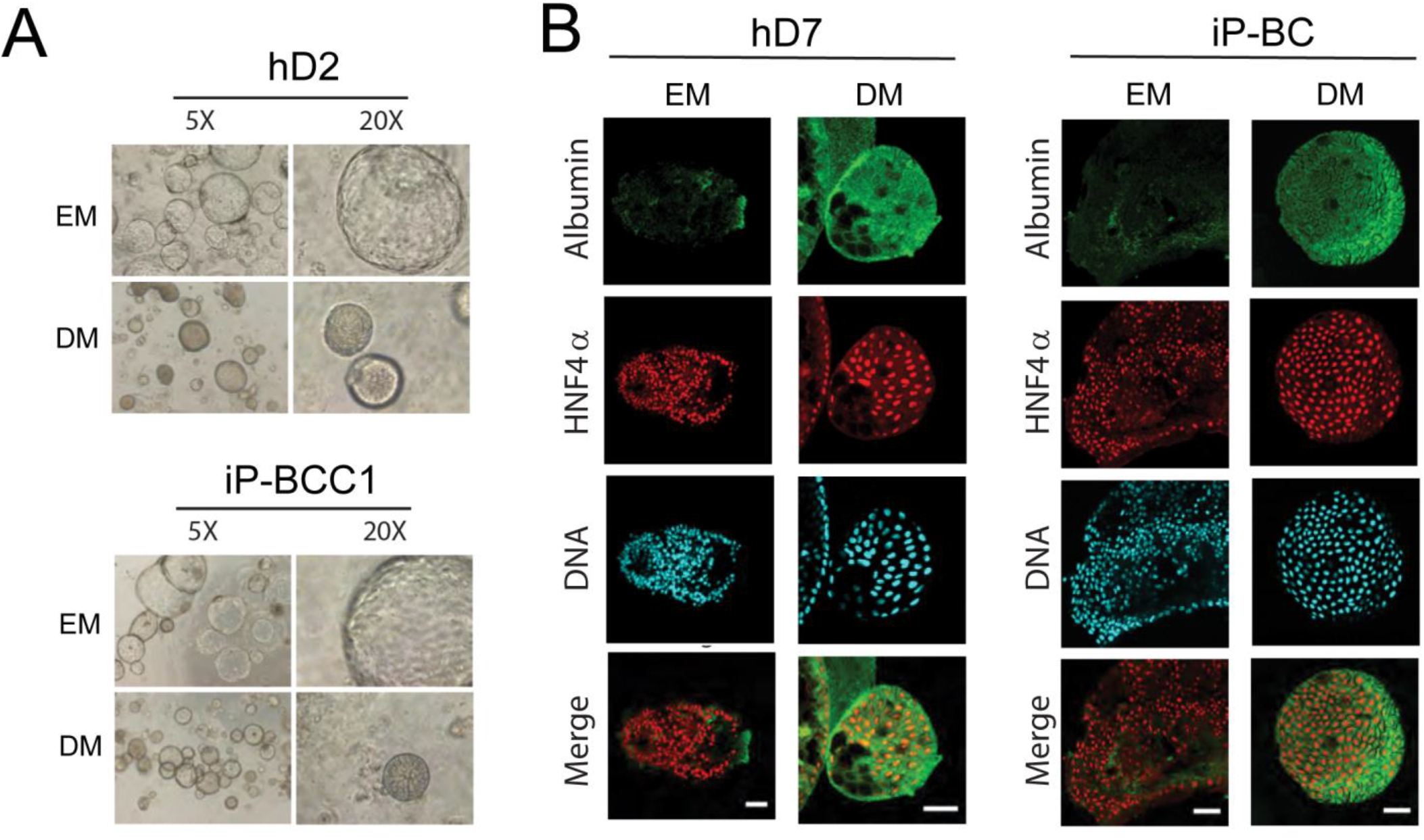
iP derived and hD derived organoids have comparable differentiation potential. (A) Phase contrast images (5X and 20X) magnification of organoid cultures seeded from hDs and iPs show comparable morphological changes in the organization upon differentiation of organoid cultures. (B) Immunofluorescent staining indicates comparable expression of the hepatocyte marker ALB (green) and HNF4α (red) in hD and iP liver organoids in expansion media (EM) and after 7 days of culture in differentiation media (DM). Nuclei are counterstained with Hoechst 33342 (cyan). Scale bars represent 50 μm.

**Table S1.** Characteristics of patients included in the study

| Patients | Age | Sex | Diagnosis | Treatment (Y/N) | HBsAg (P/N) | HBcAg (P/N) | HBeAg (P/N) | anti HBsAg | anti HBcAg | anti HBeAg | DNA |
| --- | --- | --- | --- | --- | --- | --- | --- | --- | --- | --- | --- |
| iD | 44 | F | DONOR, cleared HBV | N | N | NA | NA | P | P | NA | NA |
| iP-BC | 49 | M | Liver cirrhosis based on chronic HBV infection and portal hypertension | Y | P | NA | NA | P | P | NA | <20 |
| iP-BCC | 65 | M | HBV liver cirrhosis with HCC | Y | P | NA | NA | N | P | NA | NA |
| iP-BCC2 | 54 | M | Chronic liver cirrhosis along with HCC | Y | P | NA | NA | N | P | NA | <20 |
| iP-BCC3 | 68 | M | Chronic HBV liver cirrhosis with HCC, vital tumor at the moment of transplantation | Y | P | NA | NA | N | P | NA | <20 |
| iP-BCC4 | 52 | M | HBV liver cirrhosis with multiple HCC | Y | P | NA | N | N | P | P | <20 |
| iP-BFA | 54 | F | Acute liver failure with Crohn's disease | N | P | NA | N | N | P | P | <20 |
| iP-BFA2 | 62 | M | Acute liver failure based on HBV | N | P | NA | NA | P | P | NA | NA |
| iP-BDFA | 28 | M | Acute liver failure based on HBV and co-infected with HDV | N | P | NA | N | NA | P | P | 1180 |
| iP-B | 60 | F | Acute or chronic liver failure based on HBV infection | Y | P | NA | NA | N | P | NA | 1080 |
| iP-BDCC | 59 | M | HCC, HBV co-infected with HDV | Y | P | NA | NA | N | P | P | <20 |

**Table S2.**
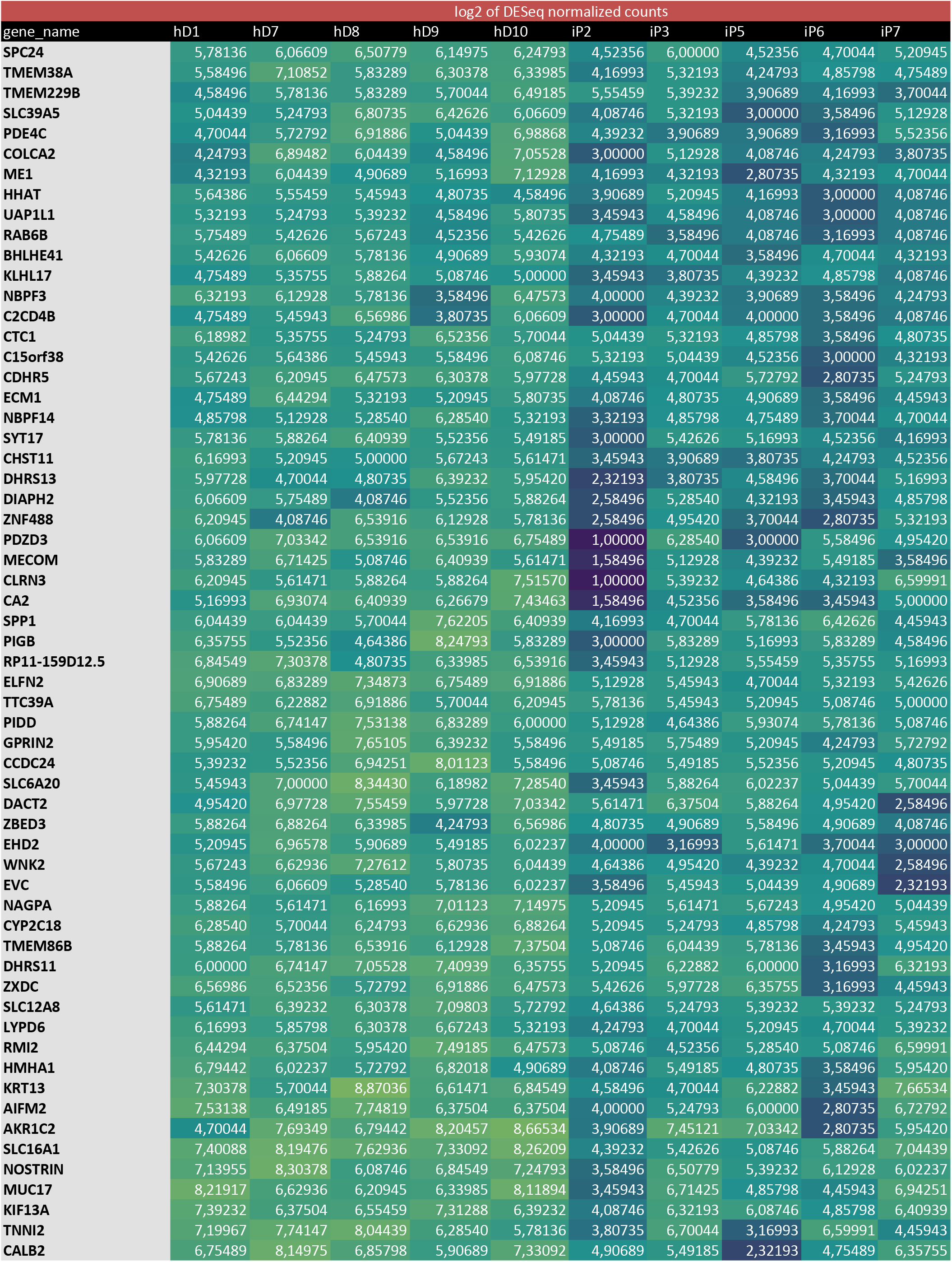
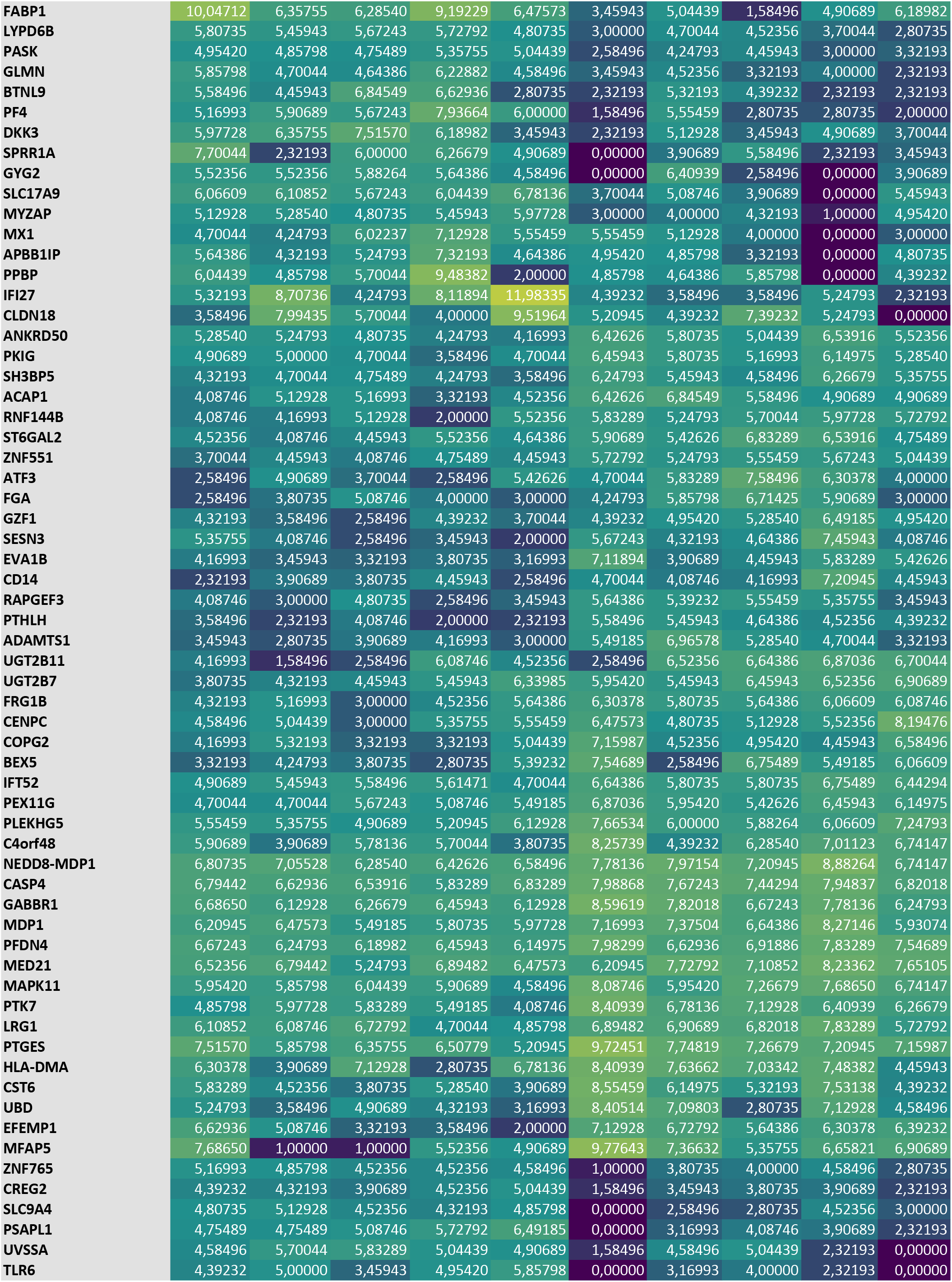
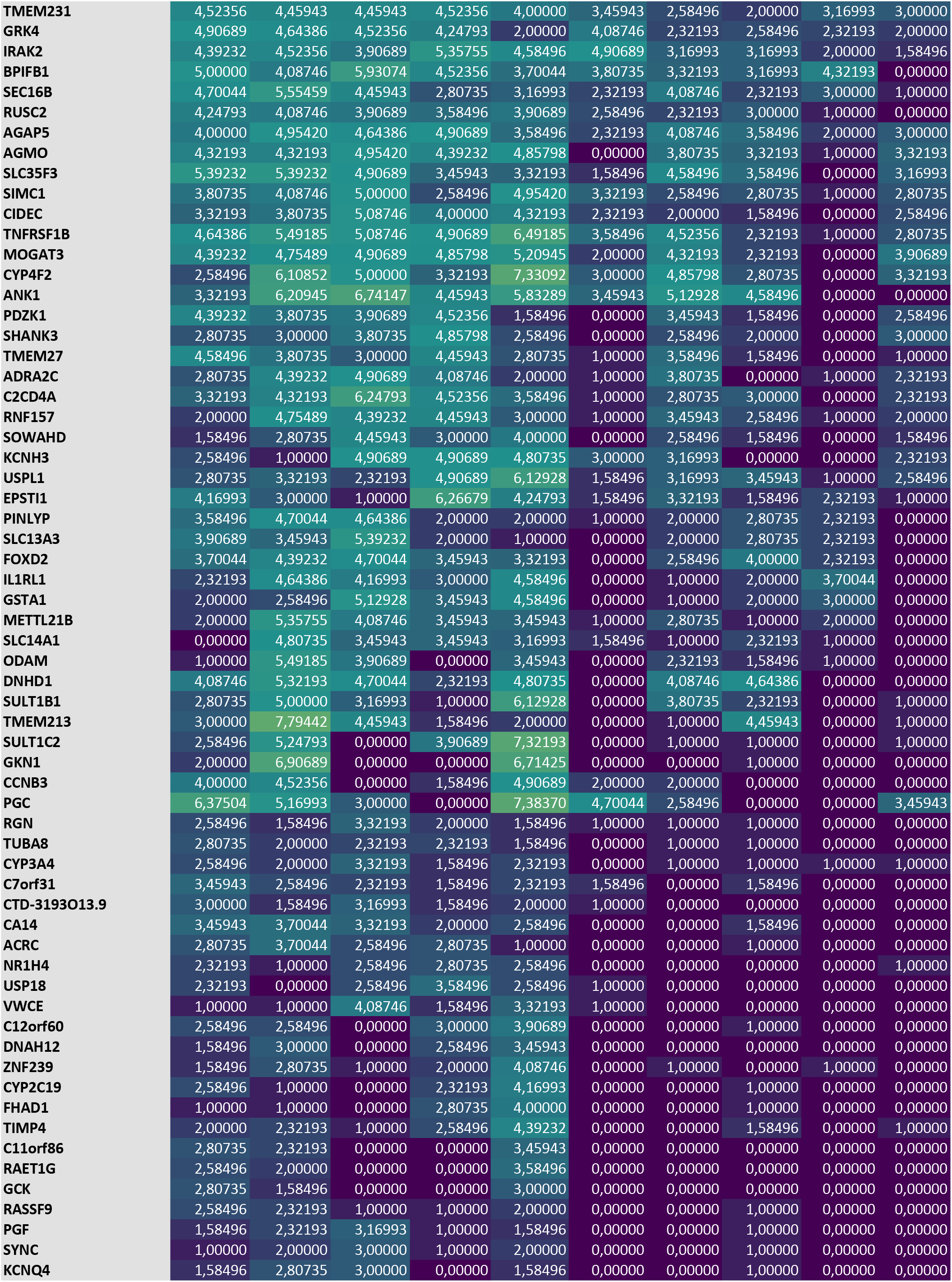
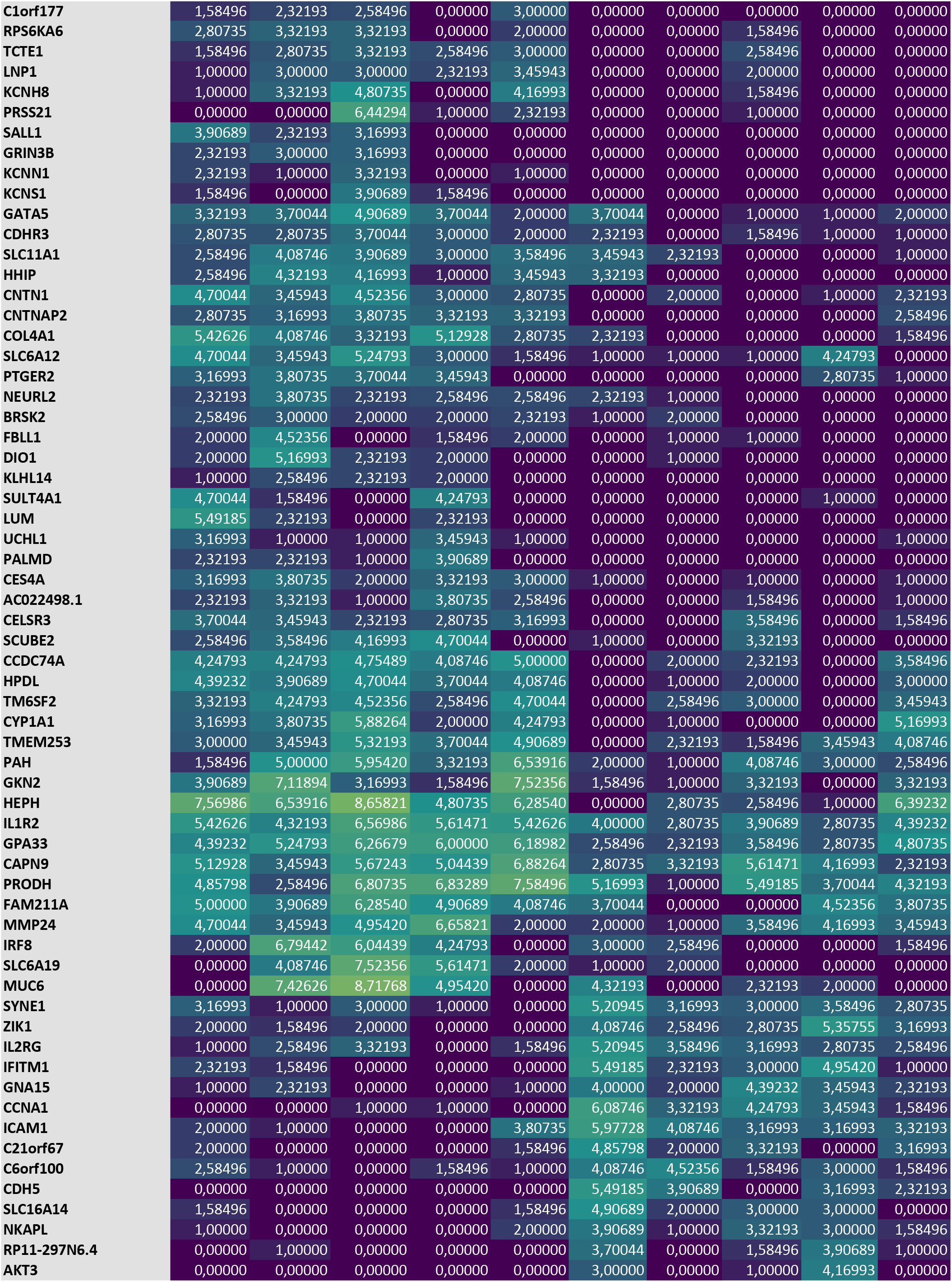
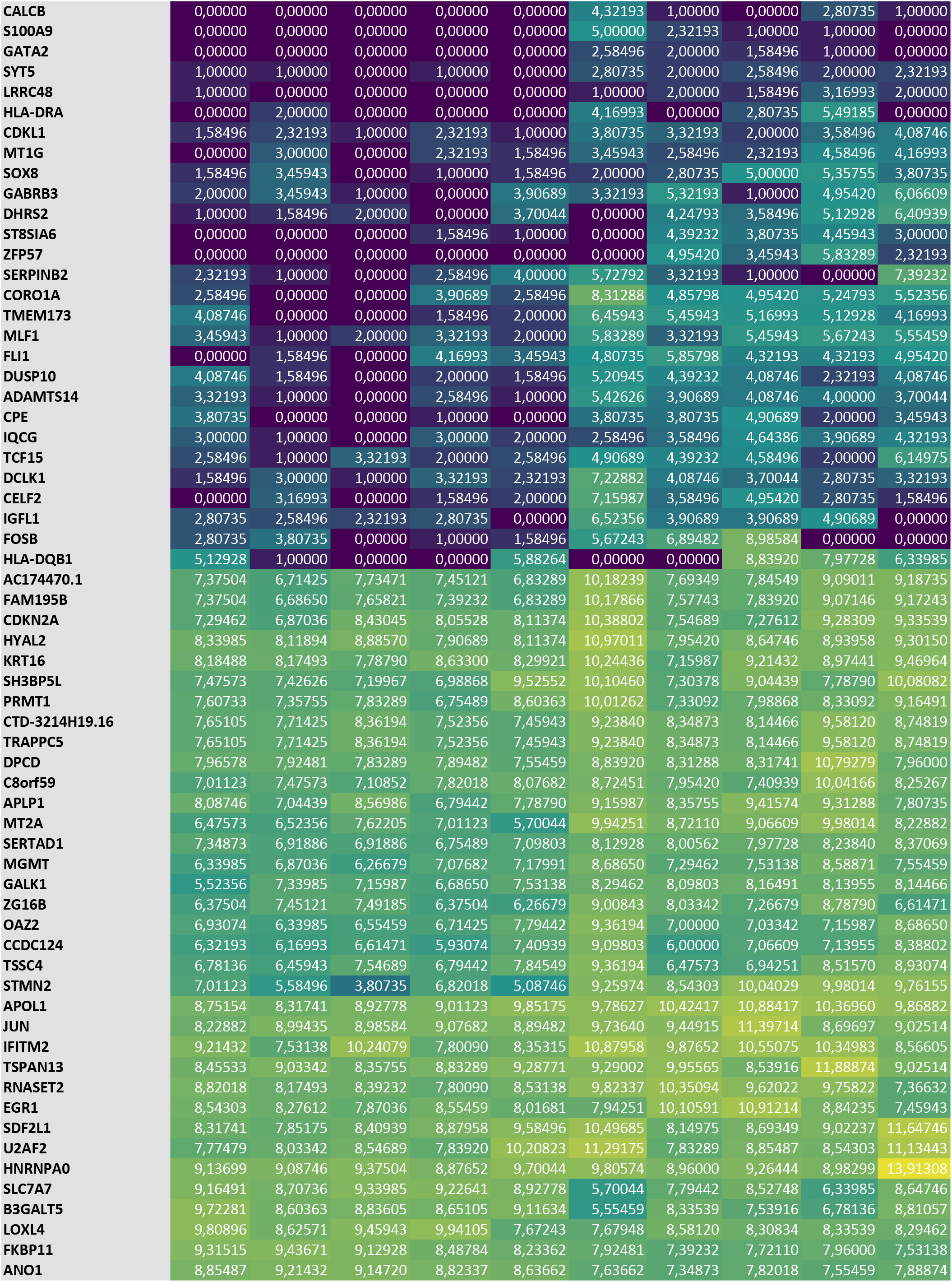
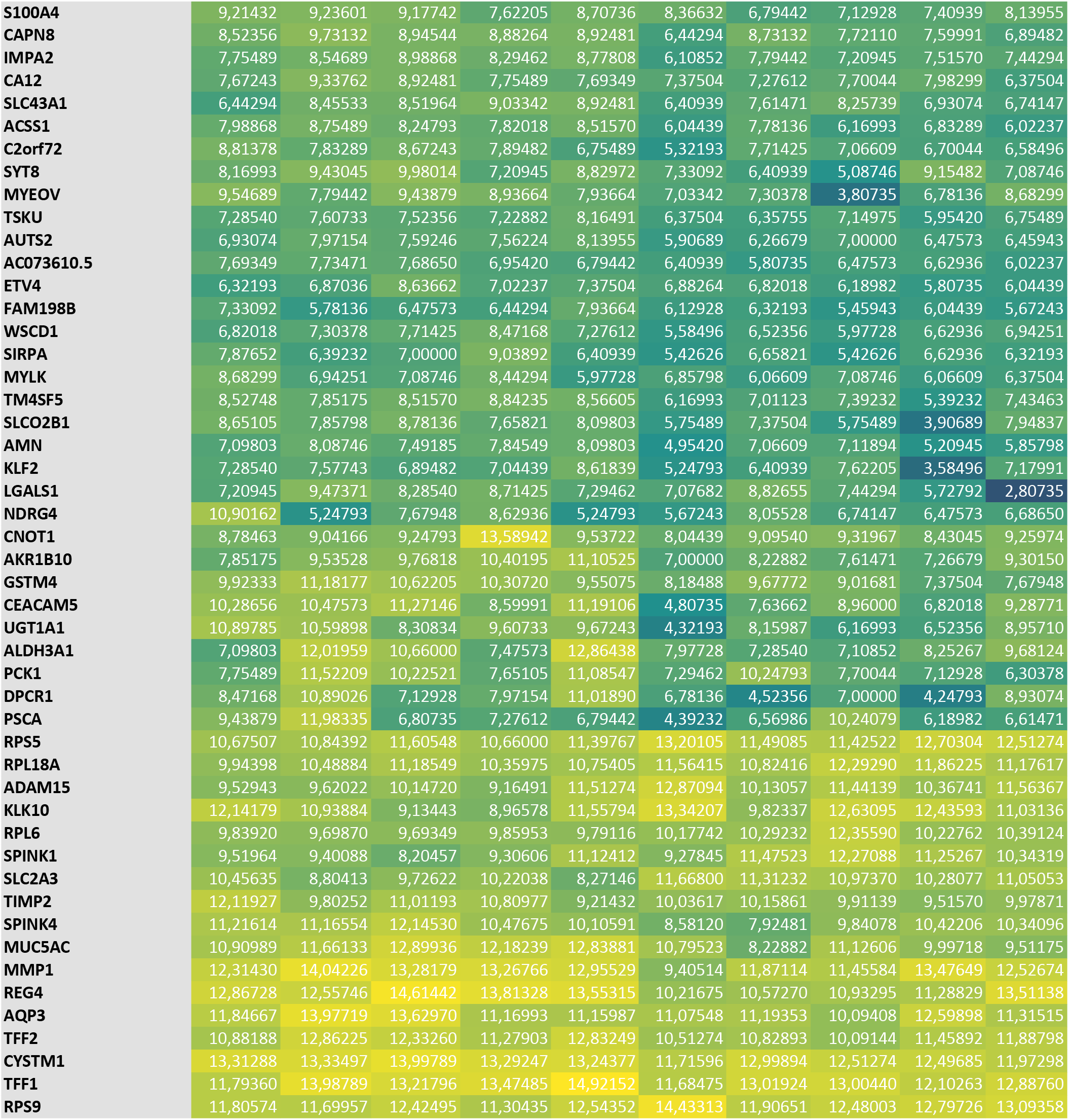
Gene expression signature

**Table S3:**
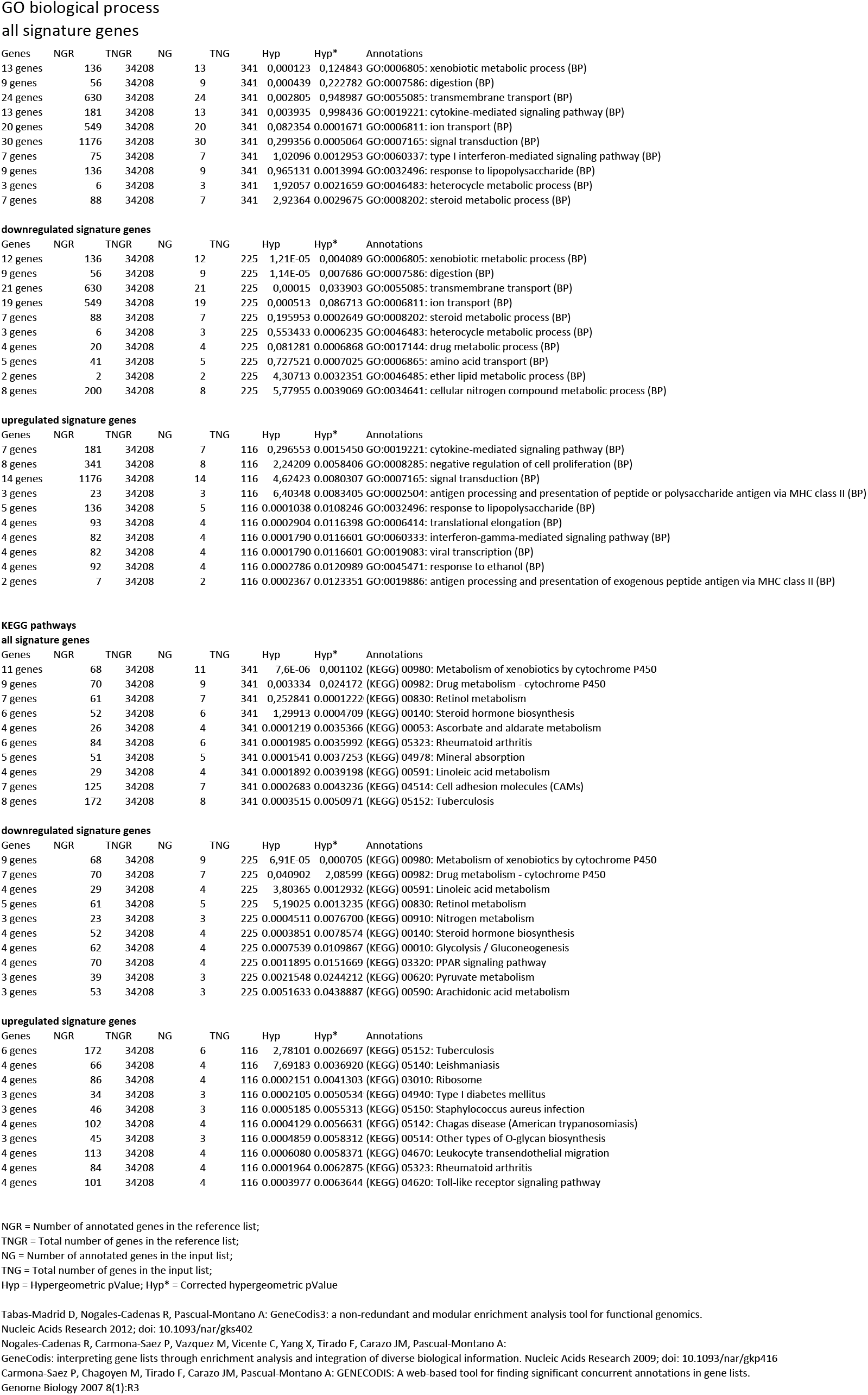
Genecodis analysis

